# Molecular Interactions of Viral Insulin/IGF-like Peptides with Zebrafish Receptors

**DOI:** 10.64898/2026.03.13.711694

**Authors:** Lev Levintov, Harish Vashisth

## Abstract

Signaling through the insulin receptor (IR) and the type 1 insulin-like growth factor receptor (IGF1R) is modulated by secreted hormones and growth factor ligands (e.g. insulin and insulin-like growth factor 1, IGF1). Impaired signaling in these receptors often leads to diabetes and oncogenic diseases. The discovery of entirely novel viral insulin/IGF-like peptides (VILPs) that can stimulate receptors from the insulin family has raised questions about their structures and binding modes to receptors. These peptides exist in a single-chain (sc) or a double-chain (dc) configuration with folds likely similar to IGF1 and insulin, respectively. The interactions of VILPs with the human receptors are beginning to be mapped but little is known about their interactions with the receptors in fish–the host organism for viruses known to carry these peptide sequences. We have previously reported [Chuard et al., Cell Rep. 2025 44(8):116149] structural models of several VILPs from the Iridoviridae virus family bound to their cognate receptors in Zebrafish (Zeb). In this work, we conducted all-atom molecular dynamics (MD) simulations of these peptides and their receptor-bound complexes along with free energy calculations to assess the energetic contributions of VILP residues for their binding to Zebrafish receptors. Most of the observed Zeb insulin/Zeb µIR and Zeb IGF1/Zeb µIGF1R site 1 interactions are consistent with previously known interactions of human peptides with their receptors, highlighting similarities in their binding modes. However, we also report some non-conserved residues in VILPs that establish significant and unique interactions with residues in Zeb receptors. Furthermore, we identified residues in each VILP which can be potentially mutated into conserved insulin/IGF1 residues to possibly enhance the binding affinity of these peptides.

## 1 INTRODUCTION

Insulin (Ins) and insulin-like growth factor 1 (IGF1) play essential roles in regulating a wide range of biological processes in cells, including metabolism, growth, and development. ^1,2^ Specifically, Ins primarily functions as a regulator of glucose homeostasis, ^1,3,4^ whereas IGF1 mainly promotes cellular growth, proliferation, and differentiation. ^2,5–7^ These peptide ligands exert their functions by binding to and activating their cognate receptors, the insulin receptor (IR) and the type 1 insulin-like growth factor receptor (IGF1R), both of which belong to the receptor tyrosine kinase (RTK) superfamily. ^7,8^ Consequently, dysregulation of IR/IGF1R leads to various diseases, most notably diabetes and cancer. ^9–12^ Therefore, significant effort has been devoted to developing potent and fast-acting insulin/IGF1 analogs with improved affinities toward receptors of the RTK superfamily^13–23^

In recent years, several organisms and pathogens have been discovered to employ molecular mimicry as a key survival strategy, enabling them to express host-like or target-like peptides and proteins. ^24–30^ For example, viruses have developed a wide range of survival mechanisms to manipulate host cell machinery by generating molecules that resemble host proteins. ^27,31–35^ Among these, viruses that infect fish have evolved viral insulin/IGF-like peptides (VILPs), ^27,36–39^ which resemble members of the insulin family. Specifically, VILPs exhibit hormone-like activity and can bind to IR and IGF1R with similar affinities. ^37^ Furthermore, these peptides have been hypothesized to adopt folds similar to those of human hormonal peptides, suggesting their potential as templates for designing novel therapeutics. ^37^

Since their discovery, several VILPs from viruses of the *Iridoviridae* family, which infect various species of fish, have been chemically synthesized. ^37–41^ These VILPs include single-chain (sc) peptides analogous to IGF1^37,41^ or double-chain (dc) peptides similar to insulin, generated by cleaving the C-domain in scVILPs (Figure S1). ^40^ Both dcVILPs and scVILPs were shown to stimulate IR and IGF1R activity *in vitro* and *in vivo*, although all of these peptides bind with higher affinities to IGF1R than to IR. ^37,39,40^ Despite these advances, only one experimental structure of a VILP bound to human IGF1R has been reported to date. ^38^ Furthermore, because of the importance of Ins/IGF1 signaling in human diseases, research on these peptides and their analogs has primarily focused on human biological systems, ^27^ resulting in an abundance of structural data for the human IR and IGF1R. ^42–51^ However, our knowledge of the binding modes of these peptides in their host organisms (e.g., fish) remains very limited. ^27,36^

Molecular modeling and molecular dynamics (MD) simulations have been successfully applied to reveal mechanistic details of the binding modes of VILPs and other insulin-like peptides to receptors from various organisms. ^48,52–64^ In a previous study, ^63^ we have successfully developed structural models of several dcVILPs derived from Grouper Iridovirus (GIV), Singapore Grouper Iridovirus (SGIV), and Lymphocystis disease virus 1 (LCDV1). We established the binding mode of each insulin-like peptide to the primary binding site of human IR, IGF1R, and hybrid IR/IGF1R and identified favorable interactions formed between these peptides and the receptors. ^63^ Furthermore, we proposed insulin analogs with enhanced interactions with the human receptors. ^63^

In this work, we report results from all-atom MD simulations of two new VILPs derived from GIV in double-chain (GIV-dcVILP) or single-chain (GIV-scVILP) configurations in their unbound states as well as their bound states to Zebrafish receptors. We also carried out free energy calculations to probe residues involved in their interactions with the receptors. Additionally, we conducted MD simulations of the structural models of native Zebrafish insulin (Zeb Ins) and Zebrafish IGF1 (Zeb IGF1) in their unbound forms and Zeb IR/IGF1R bound states ^36^ to compare against the dynamics of VILPs. The binding free energy contribution of each residue of Zeb Ins and GIV-dcVILP, Zeb IGF1 and GIV-scVILP with the receptors was further compared to identify favorable substitutions in each peptide that could potentially enhance the receptor binding affinities of insulin/IGF1 analogs.

## 2 METHODS

### 2.1 Structural models of peptides

The atomic coordinates of peptides were taken from their structural models reported in our previous work. ^36^ Briefly, we used MODELLERv10.5^65^ to model the tertiary structures of GIV-dcVILP and Zebrafish Ins (UniProtKB: O73727) based on the human Ins (hIns) structure (PDB code: 6PXV), ^43^ and GIV-scVILP and Zebrafish IGF1 (UniProtKB: Q90VV9) based on the human IGF1 (hIGF1) structure (PDB code: 6PYH). ^66^ We preserved all disulfide bonds in each peptide and the double-chain configuration of dc peptides by employing the multi-chain modeling algorithm in MODELLER. We generated 200 models of each structure and selected the best model based on the lowest discrete optimized protein energy (DOPE) score. ^67^ We used the online portal PROPKA ^68^ to assign protonation states to the sidechains (at a pH value of 7.0) of ionizable residues across all selected structures. Each peptide structure was then solvated with the OPC ^69^ water molecules and ionized with Na+ and Cl− ions at a salt concentration of 150 mM. The total number of atoms in various systems ranged between ∼24,000 and ∼28,000 atoms (Table S1).

### 2.2 Structural models of peptide/receptor complexes

To construct structural models of peptide/receptor complexes, we used the atomic coordinates of truncated Zebrafish IR (Zeb *µ*IR) and of Zebrafish IGF1R (Zeb *µ*IGF1R) from structural models reported in our previous work. ^36^ These structures were modeled using MODELLERv10.5^65^ based on the primary binding site extracted from the full-length human IR ectodomain (PDB code: 6PXV) ^43^ and the human IGF1R ectodomain (PDB: 6PYH), ^66^ respectively. We superimposed the L1 domain and the *α* CT peptide from the Zebrafish IR and IGF1R on the human *µ*IR (PDB: 6PXV) and *µ*IGF1R (PDB: 6PYH), respectively. We further superimposed peptide ligands (VILPs and Zeb Ins/IGF1) on the human Ins/IGF1 initially present in the human IR/IGF1R structures to obtain a structural complex of each ligand using the PyMOL ^70^ and Visual Molecular Dynamics (VMD) ^71^ software tools. Before conducting MD simulations, we used the online portal PROPKA ^68^ to assign protonation states (at a pH value of 7.0) to the sidechains of ionizable residues across all receptor structures. Each peptide/receptor complex was then solvated with the OPC ^69^ water molecules and ionized with Na+ and Cl− ions at a salt concentration of 150 mM. The total number of atoms in these systems ranged between ∼57,000 and ∼63,000 atoms (Table S1).

### 2.3 Molecular dynamics simulations

Each system (unbound peptides and peptide/receptor complexes) was energy-minimized via the steepest descent minimization for 1000 steps followed by 1000 steps of conjugate gradient minimization (Table S1). This was followed by a brief equilibration simulation for 0.5 ns in the NPT ensemble to equilibrate the simulation domain. Three independent MD simulations (each 500 ns long) of each unbound peptide system were performed (Table S1). For peptide/receptor complexes, we performed two independent MD simulations, each 2000 ns long (Table S1). We conducted all simulations in the NPT ensemble with a 2-fs timestep. The temperature and pressure were maintained at 300 K and 1 atm using the Langevin thermostat and the Berendsen barostat, respectively. We used a non-bonded cut-off of 12 Å with periodic boundary conditions applied across all MD simulations. The electrostatic interactions were computed using the particle mesh Ewald method. All MD simulations were conducted using the Amber ^72^ software combined with the recent ff*19SB* protein force field. ^73^ The force field parameters for solvent molecules were based on the OPC ^74^ water model which is recommended for use with the ff*19SB* protein force field. ^73^ The Li-Merz force field parameters for monovalent ions were used. ^69^ The analyses of all trajectories were carried out using the CPPTRAJ ^75^ toolkit in Amber22 package and using the VMD ^71^ software.

### 2.4 Binding free energy (ΔG_bind_) calculations

We computed the binding free energy (ΔG_bind_) of each peptide to the corresponding Zebrafish receptor as well as the interaction energy between the amino acids in each peptide with the amino acids in the Zebrafish receptors using the Molecular Mechanics/Generalized Born Surface Area (MM/GBSA) ^76–78^ approach, using a Python script (MMPBSA.py). ^79^ The binding free energy is made up of several components:

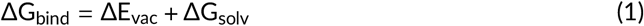

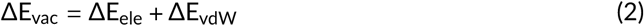

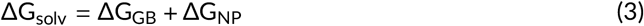

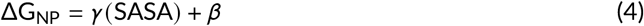

where ΔE_vac_ is the energy in vacuum (gas phase) which consists of ΔE_ele_ and ΔE_vdW_ which correspond to the electrostatic and van der Waals energies, respectively. ΔG_solv_ is the solvation free energy consisting of the polar solvation energy (ΔG_GB_) and the non-polar solvation energy (ΔG_NP_).

We calculated ΔG_NP_ by using the solvent-accessible surface area (SASA) with the surface tension of the solvent (*γ*) as 2.9 kcal Å^−2^ and the fitting parameter (*β*) as 0.0.

## 3 RESULTS

### 3.1 Conformational stability of unbound peptides

We compared the sequences of Zeb IGF1, GIV-scVILP, Zeb Ins, and GIV-dcVILP ^36^ against hIns and hIGF1. The highest sequence identity was among Zeb IGF1 and hIGF1 (∼86%), followed by Zeb Ins and hIns (∼71%), while GIV-VILPs exhibited ∼33% to ∼42% sequence identity relative to the corresponding human ligands (Figure S1). The modeled single-chain peptides retained three canonical helices in the A-and B-domains, as well as three conserved disulfide bonds present in hIGF1 (Figure 1A). Similarly, the modeled double-chain peptides also preserved three canonical helices in the A-chain and B-chain along with three conserved disulfide bonds in hIns (Figure 1B). These disulfide bonds are essential for maintaining the native three-dimensional fold of Ins and IGF1, suggesting that they likely serve the same role in VILPs. Although Zeb IGF1 closely resembled hIGF1 in its overall fold, GIV-scVILP displayed noticeable structural differences, particularly within the C-domain (Figure 1A). These differences are attributed to the C-domain of GIV-scVILP which is shorter by five residues in comparison to that of hIGF1 (Figure 1A). The overall folds of Zeb Ins and GIV-dcVILP were similar to hIns with only a slightly different orientation of the C-terminal residues of the B-chain (ProB27 through ArgB30) of GIV-dcVILP (Figure 1B).

**FIGURE 1.**
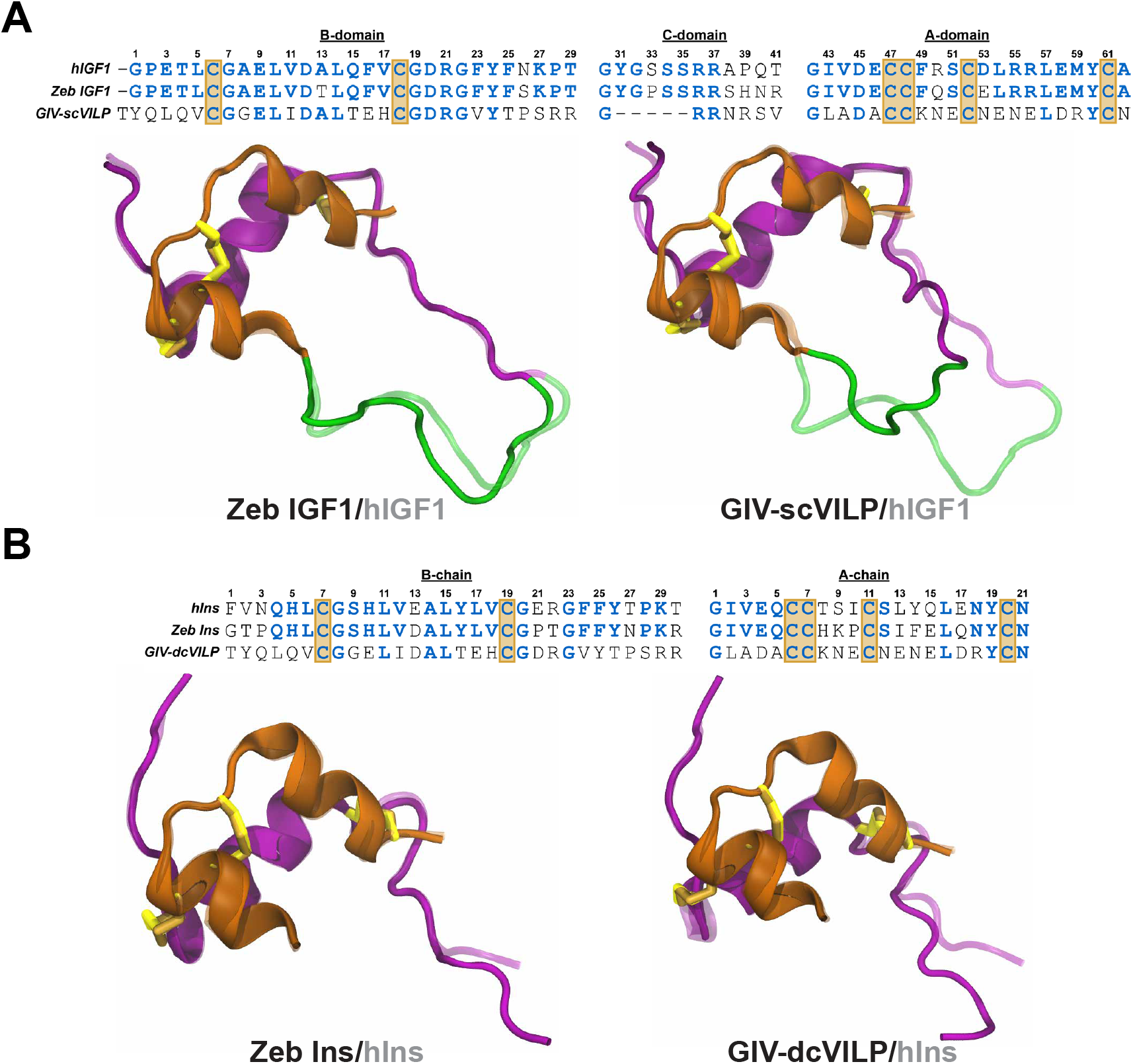
Sequence and structure alignment. The alignment of the primary sequences and the structural models (darker colors) of (A) Zeb IGF1 and GIV-scVILP on hIGF1 (PDB code: 6PYH) ^66^ and (B) Zeb Insand GIV-scVILP on hIGF1 (PDB code: 6PYH) ^66^ and (B) Zeb Ins and GIV-dcVILP on hIns (PDB code: 6PXV). ^43^ The residue numbering at the top of each sequence alignment corresponds to the hIns and hIGF1 sequences. The residues in Zeb IGF1/GIV-scVILP similar to hIGF1 and in Zeb Ins/GIV-dcVILP similar to hIns are shown in blue. The conserved cysteine residues are enclosed within golden boxes. The A-chain and the B-chain of the double-chain peptides are shown in orange and magenta, respectively; the A-domain, B-domain, and C-domain of the single-chain peptides are shown in orange, magenta, and green, respectively. The disulfide bonds in VILPs and Zeb peptides are shown as dark yellow sticks, whereas those in human peptides are depicted as light yellow sticks.

Following structural characterization, we conducted three independent all-atom MD simulations for each modeled peptide to assess their conformational flexibilities. Based on conformational data from these simulations, we identified the most populated configuration for each peptide via an RMS-based clustering analysis (Figure S2). For single-chain peptides (Zeb IGF1 and GIV-scVILP), we observed that three canonical helices from the A-domain and B-domain consistently maintained their structural integrity, indicating the stabilizing role of the conserved inter- and intra-chain disulfide bonds (Figure S2A,B). The primary distinction among the dominant clusters was the orientation of the C-domain, which exhibited notable flexibility and sampled multiple conformations relative to the core helical motifs (Figure S2A,B). The flexibility of the C-domain in each single-chain peptide was further confirmed by the backbone RMSF analysis, which showed that the C-domain residues had higher RMSF values than other residues in the entire peptide (Figure S3A).

For double-chain peptides (Zeb Ins and GIV-dcVILP), the most representative structures also demonstrated conformational stability in three helices of the A and B-chains that are held together by disulfide bonds (Figure S2C,D). A key difference between the dominant clusters of double-chain peptides was the orientation of the flexible residues at the termini of the A-chain and B-chain (Figures S2C,D and S3B). Among all modeled peptides, Zeb Ins exhibited the lowest mean heavy-atom RMSD value averaged across all MD simulations (2.82 ± 0.35 Å) signifying the most stable structure out of the modeled peptides (Figure S4). The mean RMSD value of GIV-dcVILP (4.35 ± 0.60 Å; Figure S4) was higher than the RMSD value of Zeb Ins, primarily due to the increased flexibility of the C-terminal residues of the B-chain as characterized by higher RMSF values (Figure S3B). The single-chain peptides displayed even higher mean RMSD values, with Zeb IGF1 at 5.36 ± 0.45 Å and GIV-scVILP at 5.15 ± 0.75 Å (Figure S4) primarily reflecting structural rearrangements of the C-domain observed in the most representative cluster of each unbound single-chain peptide (Figure S2A,B).

In addition to the overall heavy-atom RMSD values, we further decomposed the RMSD into contributions from individual structural segments of each peptide (Figure S4). Specifically, the RMSD values were calculated separately for the A-chain/A-domain and B-chain/B-domain to identify differences between the double-chain peptides (Zeb Ins and GIV-dcVILP) and the single-chain peptides (Zeb IGF1 and GIV-scVILP). Specifically, the A-chain of the double-chain peptides and the A-domain of the single-chain peptides exhibited similarly low mean RMSD values (Figure S4), signifying that the A-chain/A-domain remained relatively rigid across all peptides. In contrast, the B-chain of GIV-dcVILP and the B-domain of the sc peptides exhibited larger RMSD values than the corresponding A-chain/A-domain RMSD values, indicating an increased flexibility of these segments. Overall, the data on unbound peptides showed that Zeb Ins was the most stable peptide in contrast to the single-chain peptides which exhibited structural rearrangements mostly confined to their flexible C-domains.

### 3.2 Conformational stability of peptides upon receptor binding

We next characterized the dynamics of peptide–receptor complexes using all-atom MD simulations: Zeb IGF1 bound to Zeb *µ*IGF1R, GIV-scVILP bound to Zeb *µ*IGF1R, Zeb Ins bound to Zeb *µ*IR, and GIV-dcVILP bound to Zeb *µ*IR (Table S1, Figure 2). The modeled Zeb *µ*IR and Zeb *µ*IGF1R have ∼75% and ∼76% sequence identity with h*µ*IR and h*µ*IGF1R, respectively (Figure S5), suggesting similar three-dimensional folds to human receptors. In Figure 2, we show all modeled peptide/receptor complexes with the C_*α*_ atoms of the peptide residues that correspond to site 1 residues of hIns. Specifically, site 1 residues Gly7, Asp12, Leu14, Gly22, Tyr24, Gly42, and Tyr60 in hIGF1 that were found to be critical to IGF1R binding are conserved in both Zeb IGF1 and GIV-scVILP (Table S2, Figure 2A). Furthermore, site 1 residues GlyB8, LeuB11, GlyA1, TyrA19, and AsnA21 from hIns which were identified to be critical to IR binding are conserved in both Zeb Ins and GIV-dcVILP (Table S2, Figure 2B).

**FIGURE 2.**
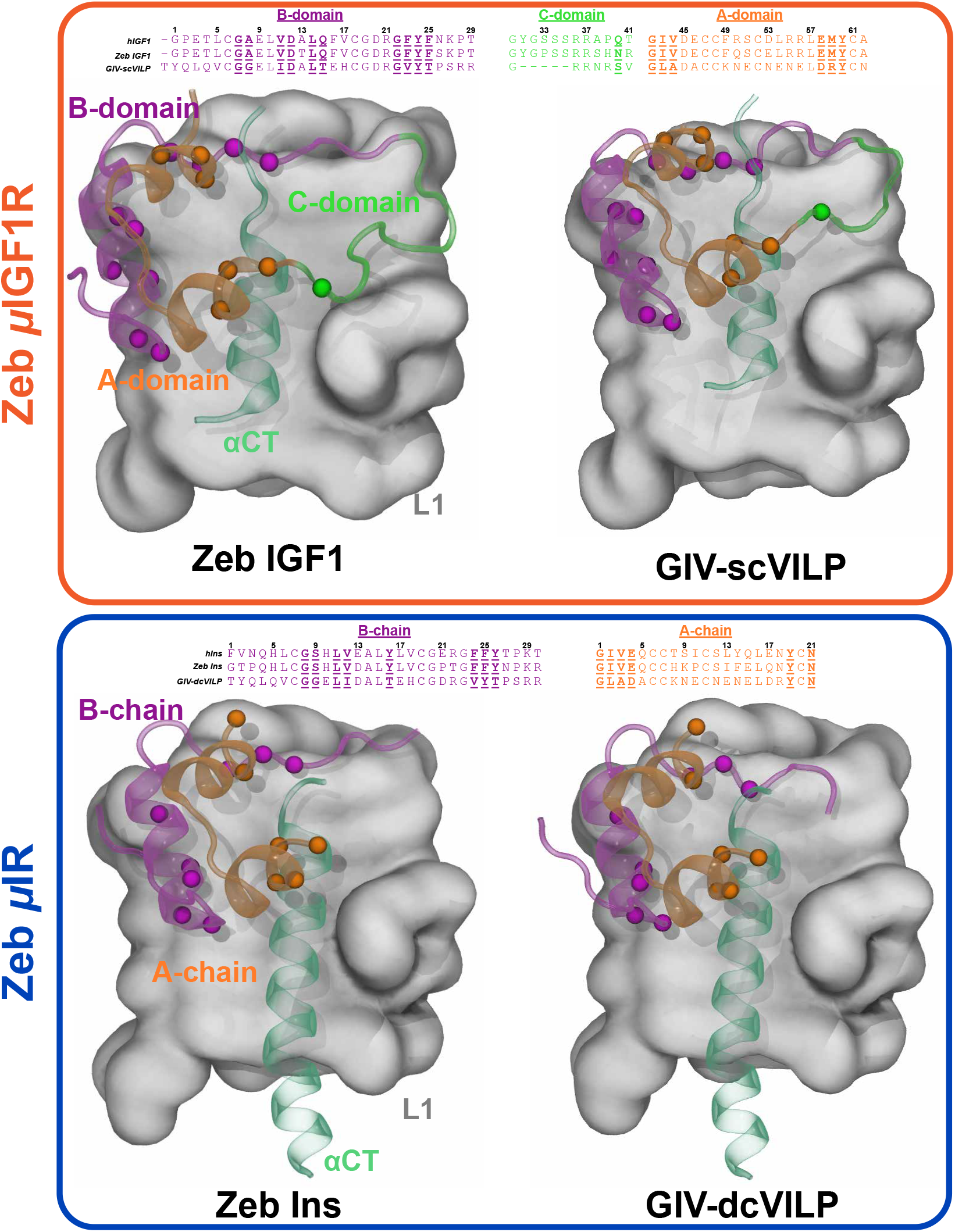
Structural models of the peptide/receptor complexes. The primary sequence alignments for each peptide are shown relative to the corresponding human variants, with site-1 residues indicated in bold underlined letters. The A-domain, B-domain, and C-domain of the single-chain peptides are shown in orange, magenta, and green, respectively; the A-chain and B-chain of the double-chain peptides are shown in orange and magenta, respectively. The C_*α*_ atoms of peptide residues corresponding to the site-1 residues in IGF1 or insulin at equivalent positions in the single-chain or double-chain peptides are depicted using spheres colored similar to the respective chain. The receptor domains are uniquely colored and labeled: L1 domain (gray surface) and *α* CT peptide (dark green).

To assess the stability of the peptide/receptor complexes, we measured the center-of-mass (COM) distance between each peptide and its corresponding receptor as well as the buried surface area (BSA) of each peptide (Figure 3). The initial COM distances for the peptide/receptor complexes ranged between 21.17 Å and 22.91 Å, and increased only marginally (by ∼1 Å) for all systems during MD simulations, except for the GIV-scVILP/*µ*IGF1R complex for which the COM distance remained mostly unchanged (Table S3, Figure 3). These values indicate that each ligand maintained contact with the corresponding receptor throughout each MD simulation. The initial BSA values for peptide/receptor complexes ranged between 1721 Å^2^ and 2689 Å^2^ and on average decreased during simulations (varying between 1360 Å^2^ and 2361 Å^2^), except for the Zeb IGF1 peptide which had an increase in BSA from 2058 Å^2^ to 2553 Å^2^ (Table S3, Figure 3). The decreases in BSA values of three peptides (Zeb Ins, GIV-dcVILP, and GIV-scVILP) were caused by structural rearrangements in residues located in the flexible regions. However, despite these structural rearrangements, significant BSA was maintained, confirming that the peptides remained stably bound to their cognate receptors. The increase in the average BSA value of Zeb IGF1 indicated a tighter contact and potentially more favorable interaction with Zeb *µ*IGF1R (Figure 3B).

**FIGURE 3.**
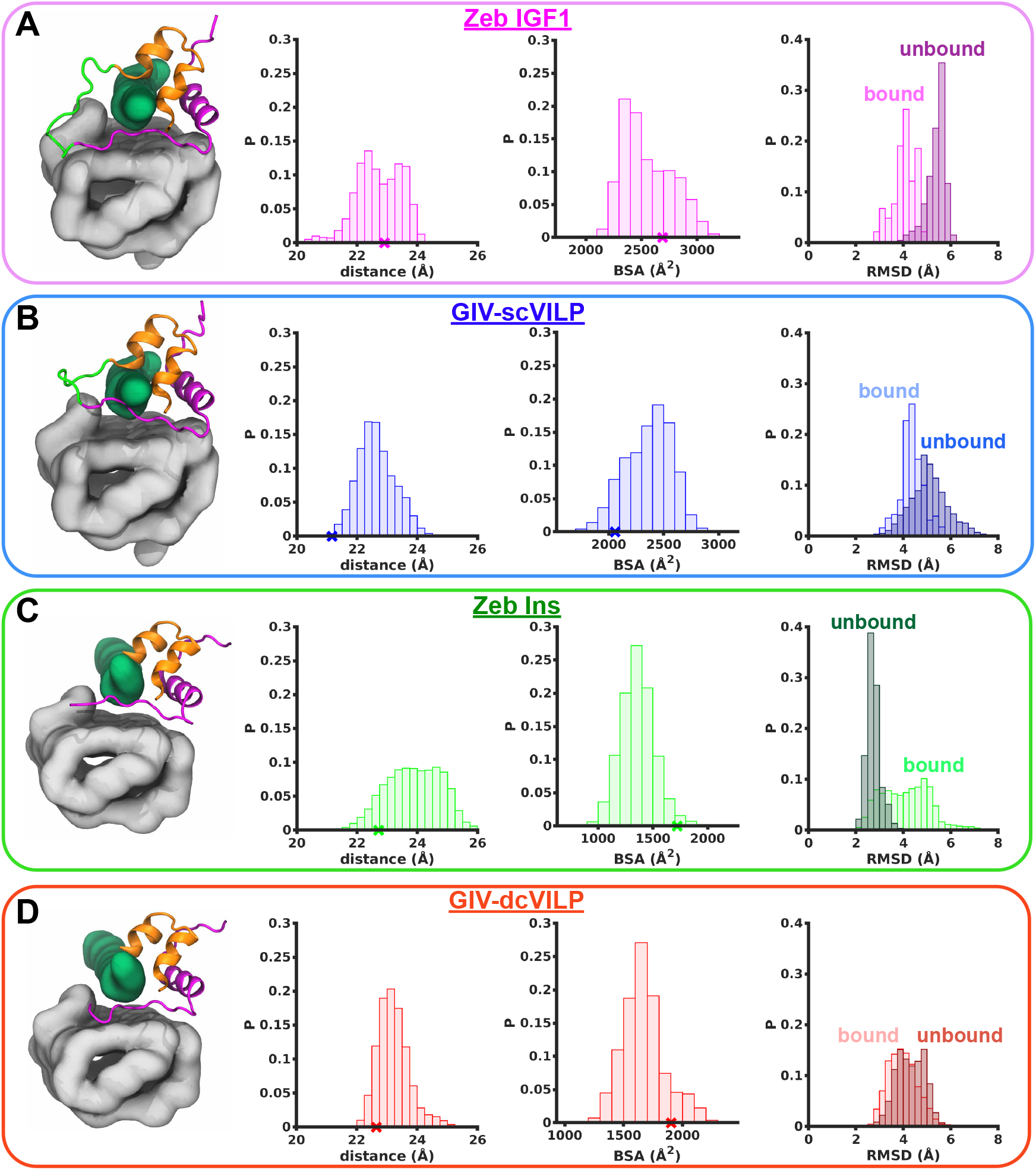
Stability metrics for each peptide/receptor complex. The side-view snapshots of each peptide/receptor complex along with the probability distributions of the COM distance between each peptide and the receptor, the BSA of each peptide, and the heavy-atom RMSD relative to initial structure for (A) Zeb IGF1, (B) GIV-scVILP, (C) Zeb Ins, and (D) GIV-dcVILP. The X symbols on the x-axis mark the initial values of the COM distance and the BSA. The RMSD of each peptide is shown in the bound (lighter colors) and unbound (darker colors) states.

Furthermore, we computed the heavy-atom RMSD of each peptide relative to the initial bound state and compared it against the RMSD of each peptide in the unbound form (Figure 3). A decrease in the mean RMSD of the bound peptide indicates stabilization of the peptide structure upon receptor binding, whereas an increase in the mean RMSD suggests enhanced structural deviations from the reference conformation upon binding. We observed a decrease in the mean RMSD values of three peptides (GIV-dcVILP, Zeb IGF1, and GIV-scVILP) relative to the mean RMSD values of the unbound peptides, except for Zeb Ins which exhibited an increase in the mean RMSD value in the bound state (Table S3, Figure 3). The RMSD distributions of the single-chain peptides (Zeb IGF1 and GIV-scVILP) in the bound and unbound states (Figure 3) revealed lower RMSD values of the bound peptides, signifying a more stable configuration. For the double-chain peptides, the RMSD distributions in the bound and unbound states either showed substantial overlap (GIV-dcVILP; Figure 3D) or displayed lower RMSD values in the unbound state (Zeb Ins; Figure 3C).

The increase in RMSD observed for Zeb Ins in the bound state relative to the unbound state was primarily attributed to enhanced flexibility of the N- and C-terminal residues of the B-chain, as indicated by the RMSF analysis (Figure S6A). This flexibility was further confirmed by the per-residue change in RMSF (ΔRMSF), where a positive value indicates an increase in the flexibility of a residue in the bound state (Figure S7). The C-terminal residues of the B-chain in the other double-chain peptide (GIV-dcVILP) were also the most flexible residues in the peptide (Figure S6A); however, compared to the unbound state, these residues exhibited reduced flexibility (Figure S7). As shown by ΔRMSF, all residues in GIV-dcVILP were more rigid in the bound state than in the unbound state (Figure S7). For the single-chain peptides, the bound-state dynamics largely resembled those observed in the unbound state, with the C-domain remaining the most flexible structural element (Figure S6B). However, in the bound state, interactions with Zeb *µ*IGF1R stabilized the C-domain in these peptides, resulting in increased rigidity relative to their unbound forms (Figure S7). Collectively, these results indicate an overall reduction in peptide flexibility upon receptor binding, likely driven by stabilizing interactions between key peptide residues and the receptor interface.

### 3.3 Thermodynamics of residue-residue interactions in the peptide/receptor interface

To characterize peptide/receptor interactions, we applied the MM/GBSA method (see Section 2.4) to compute the binding free energy (ΔG_bind_) between all atoms of each peptide with its corresponding receptor (Table S4, Figure S8). Zeb IGF1 binds to Zeb *µ*IGF1R with the lowest ΔG_bind_ value (−89.40 ± 11.76 kcal/mol), suggesting the strongest interaction among all peptides studied (Figure S8). GIV-scVILP had the second lowest ΔG_bind_ value (−73.75 ± 11.94 kcal/mol) implying a weaker interaction with Zeb *µ*IGF1R relative to its cognate ligand (Zeb IGF1). Among all peptides, Zeb Ins had the weakest interaction with its cognate Zeb *µ*IR receptor, as characterized by the highest ΔG_bind_ value (−34.65 ± 8.07; Figure S8). GIV-dcVILP had a lower ΔG_bind_ value (−52.52 ± 12.18) than the ΔG_bind_ value of Zeb Ins, signifying a more favorable interaction with Zeb *µ*IR than Zeb Ins.

We further conducted a free energy decomposition analysis on each complex using the MM/GBSA approach to identify those peptide residues that contribute most significantly to receptor binding (Figure 4). We used a cutoff of-1.5 kcal/mol based on the average per-residue ΔG_bind_ value to distinguish residues with more significant contributions to receptor binding (Figure 4). Furthermore, since no biochemical data are currently available to define the key receptor-binding residues in Zeb Ins, Zeb IGF1, or VILPs, we mapped residues corresponding to site 1 positions in hIns and hIGF1 (shown in Figure 2 and Table S2) and report their ΔG_bind_ values in Figure 4.

**FIGURE 4.**
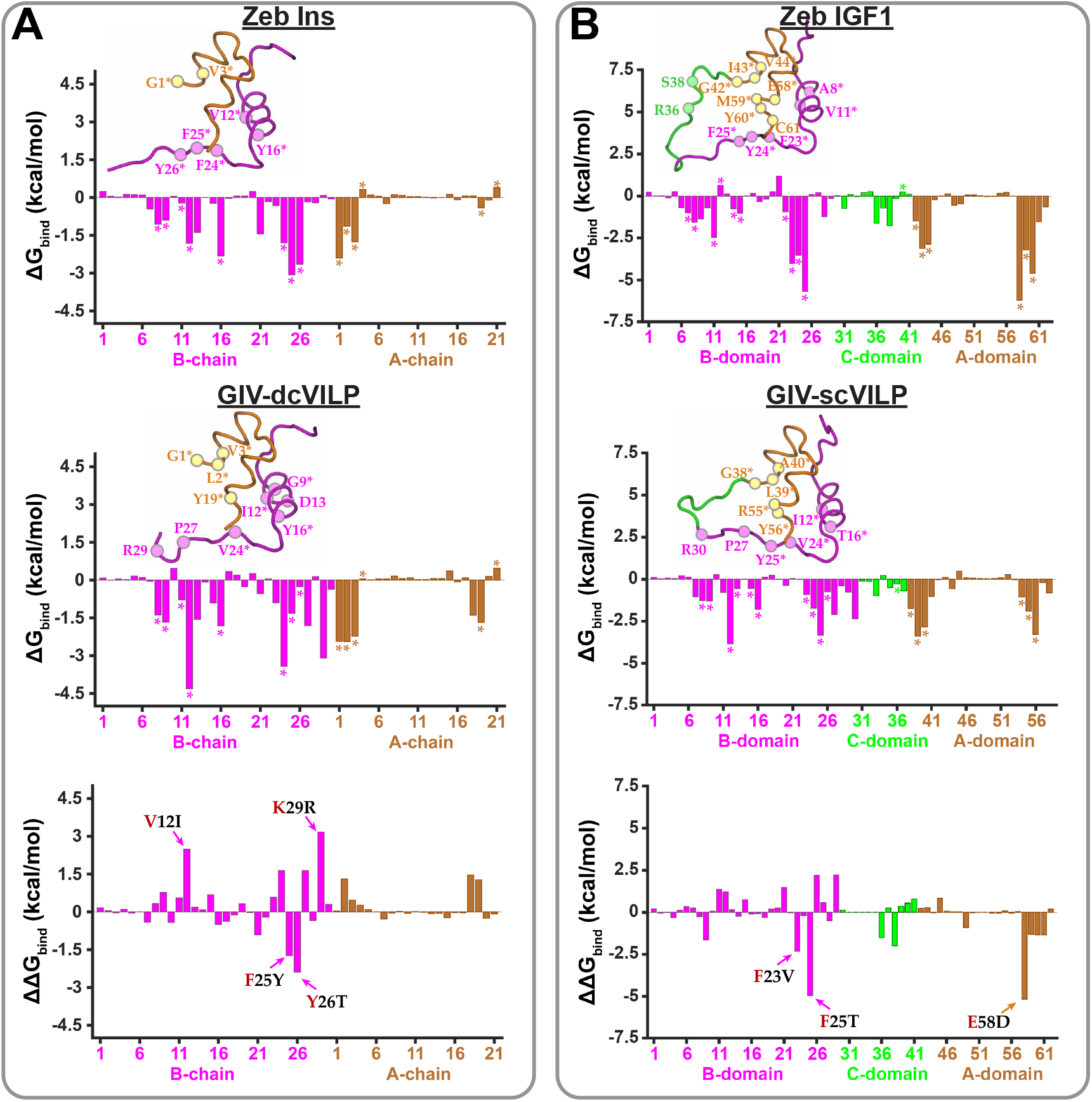
Energetics of residue-residue interactions in each peptide/receptor complex. (A) The ΔG_bind_ computed between each dc peptide residue and all residues in each receptor for two complexes (Zeb Ins/Zeb *µ*IR and GIV-dcVILP/Zeb *µ*IR) are depicted as orange bars (A-chain) and magenta bars (B-chain). The per residue energy differences (ΔΔG_bind_) between Zeb Ins and GIV-dcVILP are shown in the bottom panel. (B) Data similar to panel A are shown for two other complexes (Zeb IGF1/Zeb *µ*IGF1R and GIV-scVILP/Zeb *µ*IGF1R) with a similar color scheme for the A-domain/B-domain, while the values of the C-domain are shown as green bars. Since Zeb IGF1 and GIV-scVILP differ in sequence length, residue numbering follows the IGF1 scheme. Accordingly, the ΔΔG_bind_ values are set to zero for residues 31-35 which are absent in GIV-scVILP. The *inset* snapshots highlight peptide structures in a tube representation with the individual motifs uniquely colored (cf. Figure 1). The C_*α*_ atoms of the most critical residues based on the ΔG_bind_ analysis are shown as spheres and labeled/colored based on the respective chain. The equivalent site 1 residues to hIns (in panel A) and hIGF1 (in panel B) are marked with asterisks in the energy plots and in the *inset* snapshots (refer to Table S2 for a detailed list of site 1 residues).

Specifically, we identified residues ValB12, PheB24, PheB25, TyrB26, GlyA1, and ValA3 in Zeb Ins to be significant for binding to Zeb *µ*IR (top panel; Figure 4A). Furthermore, residues GlyB8, SerB9, IleA2, and TyrA19 also formed favorable interactions with Zeb *µ*IR, although their contributions were comparatively smaller (Table S5 and top panel in Figure 4A). All of these residues are conserved site 1 residues in hIns (Table S2), indicating that the key receptor-interacting residues are largely preserved in Zeb Ins. In GIV-dcVILP, we identified GlyB9, IleB12, AspB13, TyrB16, ValB24, ProB27, ArgB29, GlyA1, LeuA2, ValA3, and TyrA19 residues as significant residues for binding to Zeb *µ*IR (middle panel; Figure 4A). Additional residues — including GlyB8, LeuB11, TyrB25, and ThrB26 — also formed favorable interactions with Zeb *µ*IR, although their energetic contributions were comparatively smaller (Table S5 and middle panel in Figure 4A). Most of these residues correspond to site 1 positions in hIns, except for AspB13, ProB27, and ArgB29, which are not conserved in hIns (Table S2).

We further calculated per-residue differences in binding free energy using 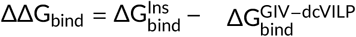 to directly compare residue-level contributions between Zeb Ins and GIV-dcVILP (bottom panel; Figure 4A). A negative ΔΔG_bind_ indicates that the corresponding GIV-dcVILP residue interacts less favorably (or not at all) with the receptor relative to Zeb Ins, whereas a positive value indicates stronger interactions for GIV-dcVILP. We observed that GIV-dcVILP had stronger interactions with Zeb *µ*IR than Zeb Ins (Table S5), although several residues in Zeb Ins exhibited stronger interactions than the corresponding GIV-dcVILP residues (F25Y and Y26T; bottom panel in Figure 4A). Specifically, PheB25 and TyrB26 in Zeb Ins were more favorable for receptor binding than TyrB25 and ThrB26 in GIV-dcVILP, respectively (Table S5 and Figure 4A).

Notably, PheB25 is a conserved hIns residue, and mutations at this position have been shown to reduce human insulin binding affinity to IR. ^80^ Thus, the binding affinity of GIV-dcVILP could be potentially enhanced by mutating TyrB25 and ThrB26 to conserved residues PheB25 and TyrB26, respectively. Additionally, we observed residues IleB12 and ArgB29 in GIV-dcVILP to have more favorable interactions with the receptor than ValB12 and LysB29 in Zeb Ins, respectively (V12I and K29R; bottom panel in Figure 4A). According to known hydrophobicity scales, ^81^ Ile and Arg are more hydrophobic than Val and Lys, respectively, which may contribute to their enhanced receptor interactions. Overall, GIV-dcVILP was a more favorable binder to Zeb *µ*IR than Zeb Ins due to a higher number of critical receptor-interacting residues (such as IleB12 and ArgB29) than Zeb Ins.

In the IGF1/Zeb *µ*IGF1R complex, we identified Ala8, Val11, Phe23, Tyr24, Phe25, Arg36, Ser38, Gly42, Ile43, Val44, Glu58, Met59, Tyr60 and Cys61 as significant residues contributing to receptor binding (top panel; Figure 4B). Additionally, Gly7, Leu14, Gln15, and Gly22 formed favorable interactions with the receptor, although their energetic contributions were comparatively smaller (Table S5 and top panel in Figure 4B). Most of these residues are equivalent to and conserved among site 1 residues in hIGF1, except for Arg36, Ser38, and Cys61 (Table S2). Thus, the receptor-interacting residues in Zeb IGF1 are largely maintained with respect to hIGF1.

In GIV-scVILP, we identified Ile12, Thr16, Val24, Tyr25, Pro27, Arg30, Gly38, Leu39, Ala40, Arg55, and Tyr56 residues as significant residues contributing to binding to Zeb *µ*IGF1R (middle panel; Figure 4B). Additional residues which formed favorable but less significant interactions with the receptor were Gly8, Gly9, Asp13, Leu15, Gly23, Thr26, and Asp54 (middle panel; Figure 4B).

With the exception of Pro27 and Arg30, these residues align with the site 1 residues in hIGF1 or Zeb IGF1, effectively reproducing the critical contact surface from hIGF1.

We further repeated the per-residue comparison using 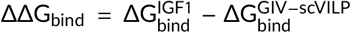 to identify residues in each single-chain peptide that had stronger interactions with the receptor (bottom panel; Figure 4B). Specifically, we observed that residues Phe23, Phe25, and Glu58 in Zeb IGF1 contribute more favorably to receptor binding than Val24, Thr26, and Asp54 in GIV-scVILP, respectively (bottom panel; Figure 4B). Phe23 and Phe25 in Zeb IGF1 are aromatic residues which are more hydrophobic than respective Val24 and Thr26 amino acids in GIV-scVILP, while Glu58 has similar hydrophobicity to Asp54, but has a longer sidechain. ^81^ Furthermore, the residues Phe23, Phe25, and Glu58 are conserved site 1 residues in hIGF1 and therefore may potentially enhance the binding affinity of GIV-scVILP to Zeb IGF1R.

To further quantify the contributions of different structural motifs of VILPs for receptor binding, we computed the cumulative energetic contributions of residues from the A-chain and B-chain of GIV-dcVILP for Zeb *µ*IR binding, and of the A-domain and B-domain of GIV-scVILP for Zeb *µ*IGF1R binding. Specifically, we summed the individual ΔG_bind_ values of residues belonging to each chain or domain reported in Figure 4. For GIV-scVILP, the B-domain residues contributed a cumulative binding energy of-22.62 kcal/mol, whereas the A-domain residues contributed only-7.72 kcal/mol, indicating a more significant contribution of the B-domain residues for Zeb *µ*IGF1R binding than the A-domain residues. This is in agreement with previous experimental results which demonstrate that the B-domain of GIV-scVILP contributes more to IGF1R binding than the A-domain. ^36^ For GIV-dcVILP, the B-chain residues contributed a cumulative binding energy of-22.28 kcal/mol, while the A-chain residues contributed-8.92 kcal/mol, highlighting a more dominant role of the B-chain in mediating interactions with Zeb *µ*IR than the A-chain.

### 3.4 Structural details of differential peptide-receptor interactions

Following the per-residue free energy decomposition analysis, we next examined the specific intermolecular interactions formed between peptides and receptors. First, we identified the residues in Zeb *µ*IR and Zeb *µ*IGF1R that adopt equivalent positions to those in h*µ*IR and h*µ*IGF1R and were previously reported to interact with hIns and hIGF1 (Table S6). ^63^ Most of these receptor residues were conserved in the Zeb homologs, with only a few substitutions observed (Table S6). Consistent with prior structural studies in human insulin and IGF1 systems, ^62,63^ the majority of reported peptide–receptor interactions were preserved in our MD simulations of Zeb homologs (Tables S7,S8), indicating conservation of the core binding interface. However, given the observed differences in binding free energy for several non-equivalent residues in the Zeb Ins/GIV-dcVILP and Zeb IGF1/GIV-scVILP peptides (Figure 4), we further investigated how these differences influence peptide interactions with each Zeb receptor (Figure 5).

**FIGURE 5.**
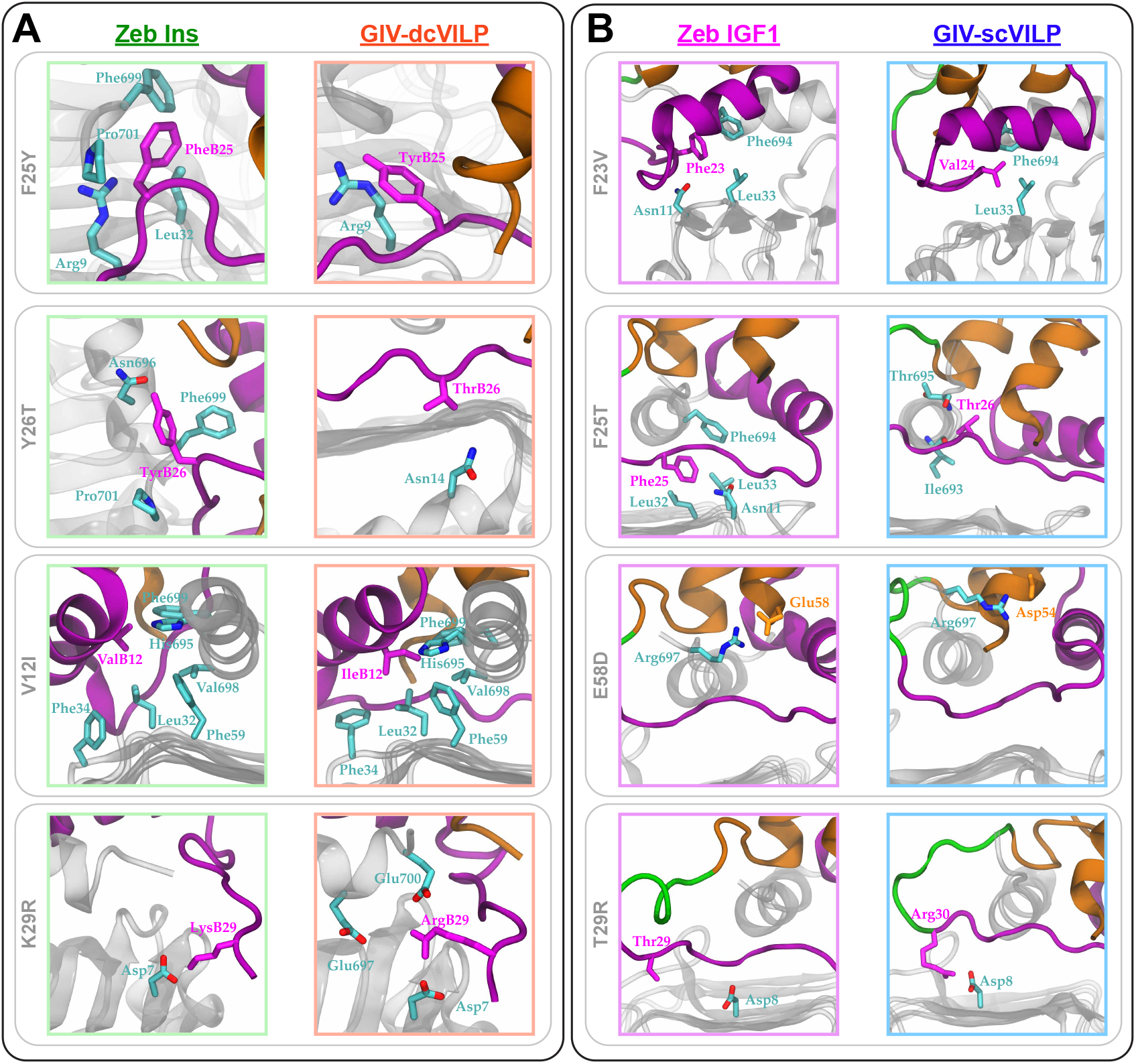
Unique inter-residue interactions between each peptide and the corresponding receptor. Shown are the inter-residue interactions of (A) Zeb *µ*IR with double-chain peptides (Zeb Ins and GIV-dcVILP) and (B) of Zeb *µ*IGF1R with single-chain peptides (Zeb IGF1 and GIV-scVILP). The sidechains of residues in peptides are depicted as magenta (B-chain) or orange (A-chain) sticks, and the sidechains of residues in each receptor are depicted as cyan sticks. The backbone of the L1 domain (gray) and *α* CT peptide (gray) are shown in a cartoon representation. cf. Figure 1 for the color scheme of peptides.

Specifically, in double-chain peptides, we first compared PheB25 in Zeb Ins to TyrB25 in GIV-dcVILP since our free energy analysis highlighted PheB25 as more energetically favorable for receptor binding than TyrB25 (F25Y; Figure 4A). We observed that PheB25 sits more deeply between the L1 domain and the *α* CT peptide than TyrB25 (F25Y; Figure 5A). As a result of this structural rearrangement, PheB25 could engage in hydrophobic interactions with Leu32 and Phe699, as well as stacking interactions with Pro701 (Figure 5A). Furthermore, the backbone atoms of PheB25 formed hydrogen-bonding interactions with Arg9 in the L1 domain (Figure 5A). In contrast, TyrB25 in GIV-dcVILP did not insert as deeply into the L1/*α* CT interface and formed only hydrogen-bonding interactions with Arg9 of the L1 domain (Figure 5A). This was due to the presence of a hydroxyl group in TyrB25, which increased its hydrophilicity in comparison to PheB25, ^81^ thereby reducing its propensity to insert more deeply into the binding site.

The structural rearrangements of PheB25 in Zeb Ins further induced rearrangements in TyrB26, which formed hydrogen bonding interactions with Asn696 (Y26T; Figure 5A). Additionally, the backbone atoms of TyrB26 formed hydrogen bonding interactions with the backbone atoms of Phe699 and Pro701 (Figure 5A). In GIV-dcVILP, ThrB26 is a shorter amino acid than TyrB26, and only interacted with Asn14 from the L1 domain (Figure 5A), thereby resulting in a lower free energy contribution than TyrB26 in Zeb Ins (Figure 4A).

Next, we examined those residues in GIV-dcVILP which were more favorable for receptor binding than residues in Zeb Ins (Figure 4). Specifically, we observed that IleB12 in GIV-dcVILP formed hydrophobic interactions with Leu32, Phe34, Phe59, Val698, and Phe699 (V12I; Figure 5A). ValB12 in Zeb Ins also formed hydrophobic interactions with these residues (Figure 5A). However, IleB12, as a more hydrophobic amino acid according to the hydrophobicity scale, ^81^ and as a slightly longer sidechain, can interact with these residues more favorably, as also captured in our free energy analysis (Figure 4).

Another unique residue pair was LysB29 in Zeb Ins and ArgB29 in GIV-dcVILP, both of which are positively-charged basic amino acids (K29R; Figure 5A). The LysB29 residue in Zeb Ins mostly interacted with Asp7 (Figure 5A), while ArgB29 in GIV-dcVILP, as a more hydrophilic amino acid, ^81^ formed salt bridging interactions with Glu697 and Glu700 (Figure 5A). Thus, a more basic ArgB29 residue could form a larger number of salt-bridging interactions with Zeb *µ*IR residues than LysB29 at the equivalent position in Zeb Ins.

We further characterized the interactions formed by the equivalent residues in the single-chain peptides (Zeb IGF1 and GIV-scVILP; Figure 5B). We observed that both Phe23 in Zeb IGF1 and Val24 in GIV-scVILP form hydrophobic interactions with Zeb *µ*IGF1R (F23V; Figure 5B). However, the free energy decomposition analysis indicated that Phe23 contributed more favorably to receptor binding than Val24 (Figure 4B). This difference was consistent with the greater hydrophobicity of Phe, ^81^ enabling stronger interactions with the hydrophobic residues within Zeb *µ*IGF1R.

Furthermore, we identified that the Phe25 residue in Zeb IGF1 was more favorable to receptor binding than the equivalent Thr26 residue in GIV-scVILP (F25T; Figure 4B). We observed that the hydrophobic Phe25 residue formed hydrophobic interactions with Gly10, Leu32, Leu33, and Phe694 (Figure 5B), while a more hydrophilic Thr26 residue in GIV-scVILP interacted with the backbone atoms of the Ile693 and Thr695 residues (Figure 5B). Thus, the deeper insertion of the more hydrophobic Phe25 residue to interact with hydrophobic residues of L1/*α* CT was energetically more favorable than the more surface-level interactions of Thr26 with Ile693 and Thr695 of the *α* CT peptide.

Another unique residue pair was Glu58 in Zeb IGF1 and Asp54 in GIV-scVILP which were both negatively-charged hydrophilic amino acids (E58D; Figure 5B). Both of these residues formed salt-bridging interactions with a positively-charged Arg697 (Figure 5B). However, the longer sidechain of Glu58 compared to that of Asp54 allowed it to adopt a more favorable conformation for effective interaction with Arg697. Lastly, Arg30 in GIV-scVILP was more energetically favorable than Thr29 in Zeb IGF1 (Figure 4B). This was due to the longer sidechain of Arg30 relative to Thr29 which resulted in stronger interactions with Asp8 from the L1 domain (T29R; Figure 5B). Overall, this comparative analysis of peptide–receptor interactions identified specific residue substitutions that could be exploited to enhance binding affinity in engineered peptide variants.

## 4 DISCUSSION

VILPs have recently emerged as a new promising class of peptides that mimic Ins and IGF1 both structurally and functionally, thereby offering new therapeutic opportunities. Experimental studies have demonstrated that VILPs can bind to IR/IGF1R in human and fish, and further stimulate autophosphorylation and downstream signaling of these receptors. ^40,41^ However, most prior experimental and computational investigations have focused primarily on VILP interactions with human receptors, ^37,38,62,63^ while comparatively little attention has been given to fish, which is the natural host of VILP-encoding viruses. ^36^ Therefore, we have conducted all-atom MD simulations of two VILPs (GIV-dcVILP and GIV-scVILP) and of two peptides from Zebrafish (Zeb Ins and Zeb IGF1) in their unbound states and in the bound states to Zebrafish *µ*IR and *µ*IGF1R (Figures 1,2).

In their unbound states, all peptides retained their characteristic structural folds, stabilized by conserved inter- and intra-chain disulfide bonds (Figure S2). For the single-chain peptides, the most significant fluctuations were localized to the C-domain, which is not stabilized by any disulfide bonds (Figure S3A), while in the double-chain peptides, the free C-terminal tail of the B-chain was the most flexible motif (Figure S3B). In the receptor bound state, peptides resembled more native like configurations than in the unbound state, with the exception of Zeb Ins which deviated relatively more in the bound state according to our RMSD analysis (Figure 3).

Consistent with these observations, the per residue ΔRMSF analysis showed that residues in GIV-dcVILP, Zeb IGF1, and GIV-scVILP became more stable upon binding to receptors (Figure S7). In Zeb Ins, the increased flexibility was primarily due to the increased residue fluctuations in the loops and terminal regions of the B-chain, whereas the helical regions of each chain remained relatively stable (Figure S7). The C-terminal tail of the B-chain in Zeb Ins contains three aromatic residues — PheB24, PheB25, and TyrB26 — which underwent conformational rearrangements to optimize interactions with residues in Zeb *µ*IR. In GIV-dcVILP, these positions are occupied by ValB24, TyrB25, and ThrB26, two of which are smaller residues relative to equivalent residues in Zeb Ins, resulting in comparatively smaller structural adjustments upon Zeb *µ*IR binding.

We further assessed the energetic contribution of each residue in each peptide for their binding to the receptor (Figure 4). We observed that Zeb Ins had ∼71% sequence conservation relative to hIns (Figures 1B, S1A), with all site 1 residues conserved (Table S2). In addition, the L1 domain and the *α* CT peptide of Zeb *µ*IR also had the majority of the reported site 1 residues to be conserved relative to h*µ*IR (Table S6). As a result of such sequence conservation in the peptides and *µ*IRs, we observed the majority of previously reported interactions in our MD simulations (Table S7). However, GIV-dcVILP exhibited substantially lower sequence similarity (∼33%-37%) relative to hIns and Zeb Ins, and many equivalent site 1 residues were not conserved (Figures 1B,S1A). Specifically, IleB12 in GIV-dcVILP was more energetically favorable (by ∼2.5 kcal/mol) than the conserved ValB12 residue in Zeb ins (Figure 4). Although both residues interacted with largely the same Zeb *µ*IR residues (Figure 5A), IleB12 was a more hydrophobic residue ^81^ with a slightly longer sidechain than ValB12, resulting in more favorable packing interactions with neighboring residues.

Furthermore, we did not observe a lower energetic contribution of SerB28 in GIV-dcVILP for receptor binding than ProB28 in Zeb Ins, as previously hypothesized (Figure 4A). ^39^ However, we observed the neighboring residues in GIV-dcVILP (ProB27 and ArgB29) to be more favorable than AsnB27 and LysB29 (Figure 4A). Specifically, ArgB29 could form favorable salt-bridging interactions with three negatively-charged amino acid sidechains, while conserved LysB29 residue in Zeb Ins could only form a single salt-bridging interaction with the Asp7 residue (Figure 5A). In Zeb Ins, the conserved residues PheB25 and TyrB26 were more favorable than the non-conserved residues TyrB25 and ThrB26 according to the free energy analysis (Figure 4A). This difference was attributed to greater hydrophobicities of PheB25 and TyrB26 in Zeb Ins compared to TyrB25 and ThrB26 in GIV-dcVILP, respectively, enabling more favorable interactions between Zeb Ins residues and the hydrophobic regions of Zeb *µ*IR (Figure 5A). Overall, this residue-level analysis identified specific substitution sites in each double-chain peptide that may serve as targets for rational mutagenesis to enhance receptor binding affinity.

For the single-chain peptides, Zeb IGF1 exhibited a high sequence conservation relative to hIGF1 (∼86%), whereas GIV-scVILP showed substantially lower similarity (∼42%; Figure S1B). Furthermore, the majority of the reported site 1 residues in the L1 domain and the *α* CT peptide of hIGF1R were also conserved in Zeb *µ*IGF1R (Table S6). As expected, we observed the majority of the previously reported interactions between IGF1 and *µ*IGF1R to be maintained in our MD simulations of Zeb IGF1 and Zeb *µ*IGF1R (Table S8). Therefore, we further compared the energetic contributions (Table S5) of the unique residues in Zeb IGF1 to GIV-scVILP to identify potential mutation sites in either peptide. Although no binding affinity measurements are currently available for these peptides with Zeb receptors, prior studies have shown that hIGF1 binds more strongly to hIGF1R than various double- and single-chain VILPs. ^39,40^ In this work, we also observed that Zeb IGF1 had a higher ΔG_bind_ than GIV-scVILP and a tighter contact according to the BSA analysis (Figure 3A,B), suggesting a more favorable interaction with Zeb *µ*IGF1R (Table S4; Figure S8).

This enhanced binding was primarily attributed to several residues in Zeb IGF1 that contributed more favorably to receptor interactions than their counterparts in GIV-scVILP, as well as to the longer C-domain present in Zeb IGF1 (Figure 4F). Specifically, two aromatic residues (Phe23 and Phe25) in Zeb IGF1 formed more favorable hydrophobic interactions with receptor residues than the equivalent Val24 and Thr26 residues in GIV-scVILP (Figure 5B). Additionally, Glu58 in Zeb IGF1 was more favorable than the equivalent Asp54 residue in GIV-scVILP due to a longer sidechain of Glu58 which allowed the residue to interact with Arg697 (Figure 5B). Conversely, Arg30 in GIV-scVILP contributed more favorably to binding than the equivalent Thr29 residue in Zeb IGF1, reflecting the advantage of a longer, positively charged sidechain at this position (Figures 4,5B).

GIV VILPs studied here were shown to initiate phosphorylation in Zeb IR and Zeb IGF1R as well as to stimulate the phosphatidylinositol 3-kinase (PI3K) pathway. ^36^ Furthermore, the previous study has demonstrated that the B-domains of GIV-VILPs are essential for their binding to IGF1R. ^36^ Consistent with these observations, our free-energy analysis revealed that residues from the B-domain of GIV-scVILP contributed substantially more to binding with Zeb *µ*IGF1R than residues from the A-domain. Nevertheless, the negative cumulative free-energy contribution from the A-domain residues indicates that they also participate in stabilizing the peptide–receptor complex.

Overall, a comparative energetic analysis presented in our work identified multiple residue positions in both peptides that may serve as targets for rational mutagenesis to enhance receptor binding affinity. Together with our previous modeling studies of Con-Ins peptides ^62^ and VILPs, ^63^ this work provides detailed residue-level mapping of peptide–receptor interactions and highlights candidate residue substitutions for further affinity optimization.

## 5 CONCLUSIONS

We conducted all-atom MD simulations on structural models of unbound peptides (Zeb Ins, GIV-dcVILP, Zeb IGF1, and GIV-scVILP) as well as their complexes with two receptors (Zeb *µ*IR and Zeb *µ*IGF1R). ^36^ All peptides maintained their characteristic folded conformations in both unbound and bound states. Receptor binding generally stabilized the peptides, with the exception of Zeb Ins, which exhibited enhanced flexibility in the C-terminal region of the B-chain. Across all complexes, conserved residues in Zeb Ins and GIV-dcVILP maintained key interactions observed in hIns/IR complexes, while Zeb IGF1 and GIV-scVILP reproduced major interaction patterns characteristic of hIGF1/IGF1R binding. However, non-conserved residues formed distinct, peptide-specific interactions that modulated overall binding energetics. The free energy decomposition analysis enabled us to probe those interactions that contributed most significantly to receptor binding and to identify residues that could potentially be modified to enhance affinity. Specifically, we identified several site 1 residues in Zeb Ins (e.g., PheB25 and TyrB26) and in Zeb IGF1 (e.g., Phe23, Phe25, and Glu58) which were more favorable to Zeb *µ*IR or Zeb *µ*IGF1R binding than the equivalent residues in GIV-dcVILP and GIV-scVILP, respectively. Thus, these residues in VILPs represent promising mutation targets for improving their receptor-binding properties. Overall, our findings offer novel opportunities for developing new peptide-based therapeutics for treatment of diseases related to proteins of the insulin receptor family.

## Abbreviations

IGF1R: type 1 insulin-like growth factor receptor
IR: insulin receptor
GIV-dcVILP: Grouper Iridovirus double-chain viral insulin-like peptide
GIV-scVILP: Grouper Iridovirus single-chain viral insulin-like peptide
MD: molecular dynamics

## 6 ACKNOWLEDGMENTS

We acknowledge financial support from the National Institutes of Health (R35GM138217). We also acknowledge computational support through the following resources: Premise, a central shared HPC cluster at UNH supported by the Research Computing Center; BioMade, a heterogeneous CPU/GPU cluster supported by the NSF EPSCoR award (OIA-1757371).

## 8 SUPPORTING INFORMATION

The data that support the findings of this study are available in the supplementary material of this article.

