## Supporting Information for "Molecular Interactions of Viral Insulin/IGF-like Peptides with Zebrafish Receptors"

**Table S1.** Simulation setup and details of all-atom MD simulations.

| System | System size (total atoms) | Simulation domain (Å$^3$) | Minimization (steps) | Trajectory length (ns) | Replicas |
|---|---|---|---|---|---|
| | | *Peptides* | | | |
| Zeb Ins | 26,593 | $70 \times 62 \times 59$ | 2000 | 500 | 3 |
| GIV-dcVILP | 24,741 | $72 \times 63 \times 54$ | 2000 | 500 | 3 |
| Zeb IGF1 | 27,792 | $68 \times 64 \times 62$ | 2000 | 500 | 3 |
| GIV-scVILP | 24,675 | $64 \times 61 \times 63$ | 2000 | 500 | 3 |
| | | *Complexes* | | | |
| Zeb Ins/Zeb $\mu$IR | 63,235 | $84 \times 84 \times 84$ | 2000 | 2000 | 2 |
| GIV-dcVILP/Zeb $\mu$IR | 60,003 | $84 \times 84 \times 79$ | 2000 | 2000 | 2 |
| Zeb IGF1/Zeb $\mu$IGF1R | 57,168 | $77 \times 84 \times 83$ | 2000 | 2000 | 2 |
| GIV-scVILP/Zeb $\mu$IGF1R | 57,283 | $75 \times 84 \times 84$ | 2000 | 2000 | 2 |

**Zeb Ins**: Zebrafish insulin; **GIV-dcVILP**: Grouper Iridovirus double-chain viral insulin-like peptide; **Zeb IGF1**: Zebrafish insulin-like growth factor 1; **GIV-scVILP**: Grouper Iridovirus single-chain viral insulin-like peptide; **Zeb $\mu$IR**: truncated Zebrafish insulin receptor; **Zeb $\mu$IGF1R**: truncated Zebrafish type 1 insulin-like growth factor receptor.

**Table S2.** A comparison of site 1 residues for the human peptides (hIns and hIGF1) and at equivalent positions in the studied peptides (cf. Figure 2). The residues are colored based on chains in the double-chain peptides: A-chain (orange) and B-chain (magenta). In the single-chain peptides, the residues are colored based on domains: A-domain (orange), B-domain (magenta), and C-domain (green). Underlined residues are conserved across all three double-chain or single-chain peptides. The residues marked with the # symbol are conserved across any two double-chain or single-chain peptides.

| Double-chain peptides | | | Single-chain peptides | | |
|---|---|---|---|---|---|
| **Site 1 residues of hIns** | **Site 1 residues of Zeb Ins** | **Site 1 residues of GIV-dcVILP** | **Site 1 residues of hIGF1** | **Site 1 residues of Zeb IGF1** | **Site 1 residues of GIV-scVILP** |
| GlyB8 | GlyB8 | GlyB8 | Gly7 | Gly7 | Gly8 |
| SerB9# | SerB9# | GlyB9 | Ala8# | Ala8# | Gly9 |
| LeuB11 | LeuB11 | LeuB11 | Val11# | Val11# | Ile12 |
| ValB12# | ValB12# | IleB12 | Asp12 | Asp12 | Asp13 |
| TyrB16# | TyrB16# | ThrB16 | Leu14 | Leu14 | Leu15 |
| PheB24# | PheB24# | ValB24 | Gln15# | Gln15# | Thr16 |
| PheB25# | PheB25# | TyrB25 | Gly22 | Gly22 | Gly23 |
| TyrB26# | TyrB26# | ThrB26 | Phe23# | Phe23# | Val24 |
| GlyA1 | GlyA1 | GlyA1 | Tyr24 | Tyr24 | Tyr25 |
| IleA2# | IleA2# | LeuA2 | Phe25# | Phe25# | Thr26 |
| ValA3# | ValA3# | AlaA3 | Gln40 | Asn40 | Ser36 |
| GluA4# | GluA4# | AspA4 | Gly42 | Gly42 | Gly38 |
| TyrA19 | TyrA19 | TyrA19 | Ile43# | Ile43# | Leu39 |
| AsnA21 | AsnA21 | AsnA21 | Val44# | Val44# | Ala40 |
| | | | Glu58# | Glu58# | Asp54 |
| | | | Met59# | Met59# | Arg55 |
| | | | Tyr60 | Tyr60 | Tyr56 |

**Table S3.** Conformational stability metrics for peptide/receptor complexes. Initial and average values of the center-of-mass (COM) distance and of the buried surface area (BSA), and the average heavy-atom RMSD values of the unbound and bound peptides.

| System | COM distance (Å) | | BSA (Å$^2$) | | RMSD (Å) | |
|---|---|---|---|---|---|---|
| | Initial | Average | Initial | Average | Unbound | Bound |
| Zeb Ins/Zeb $\mu$IR | 22.71 | 23.88 ± 0.84 | 1721 | 1360 ± 158 | 2.82 ± 0.35 | 4.12 ± 0.74 |
| GIV-dcVILP/Zeb $\mu$IR | 22.65 | 23.22 ± 0.53 | 1903 | 1671 ± 182 | 4.35 ± 0.60 | 3.98 ± 0.63 |
| Zeb IGF1/Zeb $\mu$IGF1R | 21.17 | 22.72 ± 0.60 | 2058 | 2553 ± 225 | 5.36 ± 0.45 | 4.04 ± 0.42 |
| GIV-scVILP/Zeb $\mu$IGF1R | 22.91 | 22.64 ± 0.58 | 2689 | 2361 ± 213 | 5.15 ± 0.75 | 4.36 ± 0.43 |

**Table S4.** The binding affinity ($\Delta G_{bind}$) computed between all-atoms of each peptide and all-atoms of the corresponding receptor.

| System | $\Delta G_{bind}$ (kcal/mol) |
|---|---|
| Zeb Ins/Zeb $\mu$IR | -34.65 ± 8.07 |
| GIV-dcVILP/Zeb $\mu$IR | -52.52 ± 12.18 |
| Zeb IGF1/Zeb $\mu$IGF1R | -89.40 ± 11.76 |
| GIV-scVILP/Zeb $\mu$IGF1R | -73.75 ± 11.94 |

**Table S5.** The binding free energy values of site 1 residues in each peptide. The residues conserved across all peptides are underlined, including in human peptides.

| Double-chain peptides | | | Single-chain peptides | | |
|---|---|---|---|---|---|
| **Residue** | **Zeb Ins** | **GIV-dcVILP** | **Residue** | **Zeb IGF1** | **GIV-scVILP** |
| **B8** | -1.05 ± 0.66 | -1.38 ± 0.62 | **B7** | -1.02 ± 0.84 | -1.28 ± 0.88 |
| **B9** | -0.90 ± 1.03 | -1.67 ± 0.62 | **B8** | -1.56 ± 0.74 | -1.30 ± 0.86 |
| **B11** | -0.22 ± 0.17 | -0.78 ± 0.50 | **B11** | -2.49 ± 0.46 | -3.85 ± 0.62 |
| **B12** | -1.82 ± 0.50 | -4.31 ± 0.81 | **B12** | 0.64 ± 0.44 | -0.57 ± 1.20 |
| **B16** | -2.32 ± 1.6 | -1.82 ± 1.29 | **B14** | -0.76 ± 0.22 | -0.54 ± 0.28 |
| **B24** | -1.79 ± 0.89 | -3.42 ± 0.79 | **B15** | -1.04 ± 0.84 | -1.79 ± 1.3 |
| **B25** | -3.06 ± 1.15 | -1.32 ± 1.10 | **B22** | -0.96 ± 0.46 | -0.92 ± 0.50 |
| **B26** | -2.64 ± 1.54 | -0.26 ± 0.67 | **B23** | -4.03 ± 0.88 | -1.72 ± 0.92 |
| **A1** | -2.40 ± 1.62 | -2.43 ± 1.46 | **B24** | -3.54 ± 0.61 | -3.34 ± 1.19 |
| **A2** | -1.14 ± 0.72 | -2.44 ± 0.80 | **B25** | -5.71 ± 0.92 | -0.75 ± 1.18 |
| **A3** | -1.76 ± 0.52 | -2.23 ± 0.76 | **C40** | 0.26 ± 0.48 | -0.28 ± 1.05 |
| **A4** | 0.32 ± 1.16 | 0.05 ± 0.91 | **A42** | -1.50 ± 0.56 | -1.74 ± 1.02 |
| **A19** | -0.41 ± 0.44 | -1.68 ± 0.68 | **A43** | -3.14 ± 0.60 | -3.41 ± 0.84 |
| **A21** | 0.40 ± 0.30 | 0.48 ± 0.29 | **A44** | -2.88 ± 0.75 | -2.84 ± 0.75 |
| | | | **A58** | -6.24 ± 1.36 | -1.06 ± 1.60 |
| | | | **A59** | -3.21 ± 0.73 | -1.90 ± 1.92 |
| | | | **A60** | -4.62 ± 0.56 | -3.28 ± 0.86 |

**Table S6.** The site 1 residues are listed for the human receptors (h$\mu$IR and h$\mu$IGF1R) and at equivalent positions in the Zebrafish receptors (Zeb $\mu$IR and Zeb $\mu$IGF1R). The underlined residues for the receptors are from the L1 domain. The residues that are not conserved in receptors are marked with the # symbol.

| **hIns** | **h$\mu$IR** | **Zeb $\mu$IR** | **hIGF1** | **h$\mu$IGF1R** | **Zeb $\mu$IGF1R** |
|---|---|---|---|---|---|
| **B8** | H710 | H695 | **B7** | E693; H697 | E686; H690 |
| **B9** | H710 | H695 | **B8** | E693; E91 | E686; E96 |
| **B11** | F714 | F699 | **B11** | H697; F701; R59 | H690; F694; R64 |
| **B12** | F39; F64; R65; H710; F714 | F34; F59; R60; H695; F699 | **B12** | R59 | R64 |
| **B16** | F39 | F34 | **B14** | F701 | F694 |
| **B24** | N15; L37; F39; F714 | N10; L32; F34; F699 | **B15** | L33; R59 | L33; R64 |
| **B25** | V715 | E700$^{\#}$ | **B22** | N11 | N11 |
| **B26** | D12; R14 | D7; R9 | **B23** | F701; R8;$^{\#}$ N11; L33 | F694; G8;$^{\#}$ N11; L38 |
| **A1** | F714; N711 | F699; N696 | **B24** | V702; P703; R704; R10$^{\#}$ | T695; P696; R697; G10$^{\#}$ |
| **A2** | F714 | F699 | **B25** | I700; D8; R10$^{\#}$ | I693; D8; G10$^{\#}$ |
| **A3** | H710; N711 | H695; N696 | **C40** | F695; N698; S699$^{\#}$ | F688; N691; A692$^{\#}$ |
| **A4** | N711 | N696 | **A42** | N698 | N691 |
| **A19** | F714 | F699 | **A43** | H697; N698 | H690; N691 |
| **A21** | N15 | N10 | **A44** | N694; N698 | N687; N691 |
| | | | **A58** | R704 | R697 |
| | | | **A59** | P703; R704 | P696; R697 |
| | | | **A60** | F701; P703 | F694; P696 |

**Table S7.** Listed are the Zeb $\mu$IR residues in contact with each site 1 residue in the A-chain (orange) and the B-chain (magenta) of Zeb Ins and GIV-dcVILP.

| Zeb Ins | Zeb $\mu$IR | GIV-dcVILP | Zeb $\mu$IR |
|---|---|---|---|
| GlyB8 | H695; F699 | GlyB8 | H695; E691 |
| SerB9 | H695; F699 | GlyB9 | H695 |
| LeuB11 | H695; F699 | LeuB11 | H695; F699 |
| ValB12 | L32; F34; F59; R60 | IleB12 | L32; F34; F59; R60 |
| TyrB16 | F34; K35 | ThrB16 | F34; R60 |
| PheB24 | N10; L32; F34; F699; P700; E701 | ValB24 | N10; L32; F34; V698; F699 |
| PheB25 | R9; N10; I11; N14; L32 | TyrB25 | R9; N10 |
| TyrB26 | R9; N10; N14; E700; P701 | ThrB26 | D7; R9 |
| GlyA1 | N692; H695; N696; F699; E700; P701 | GlyA1 | N696; F699; E700; P701 |
| IleA2 | H695; F699; E700 | LeuA2 | H695; N696; V698; F699 |
| ValA3 | H695; F699 | AlaA3 | N692; H695 |
| GlyA4 | K688; N692; H695; N696 | AspA4 | K688; N692; N696 |
| TyrA19 | E700; P701 | TyrA19 | F699; E700; P701 |
| AsnA21 | *no contacts* | AsnA21 | N10 |

**Table S8.** Listed are the Zeb $\mu$IGF1R residues in contact with each site 1 residue in the A-domain (orange), the B-domain (magenta), and the C-domain (green) of Zeb IGF1 and GIV-scVILP.

| **Zeb IGF1** | **Zeb $\mu$IGF1R** | **GIV-scVILP** | **Zeb $\mu$IGF1R** |
|---|---|---|---|
| **Gly7** | E686; N687; H690 | **Gly8** | E686; H690 |
| **Ala8** | E686; N687 | **Gly9** | R64; E686 |
| **Val11** | L33; R64; H690; F694 | **Ile12** | L33; N38; R64 |
| **Asp12** | N38; R64 | **Asp13** | N38; R64 |
| **Leu14** | F694 | **Leu15** | F694 |
| **Gln15** | D36; N38; N40 | **Thr16** | K37; N38 |
| **Gly22** | N11 | **Gly23** | N11 |
| **Phe23** | N11; L33; N38; F694 | **Val24** | N11; L33 |
| **Tyr24** | T695; P696; R697; P698 | **Tyr25** | D8; G10; N11; L32; L33 |
| **Phe25** | D8; G10; N11; L32; L33 | **Thr26** | D8 |
| **Asn40** | P696 | **Ser36** | F688; N691 |
| **Gly42** | N691; F694; P696 | **Gly38** | N691; F694 |
| **Ile43** | H690; N691; F694 | **Leu39** | H690; N691 |
| **Val44** | N687; H690; N691 | **Ala40** | N687; H690 |
| **Glu58** | R697 | **Asp54** | R697 |
| **Met59** | R696; R697 | **Arg55** | P696; P698 |
| **Tyr60** | F694 | **Tyr56** | F694; P696 |

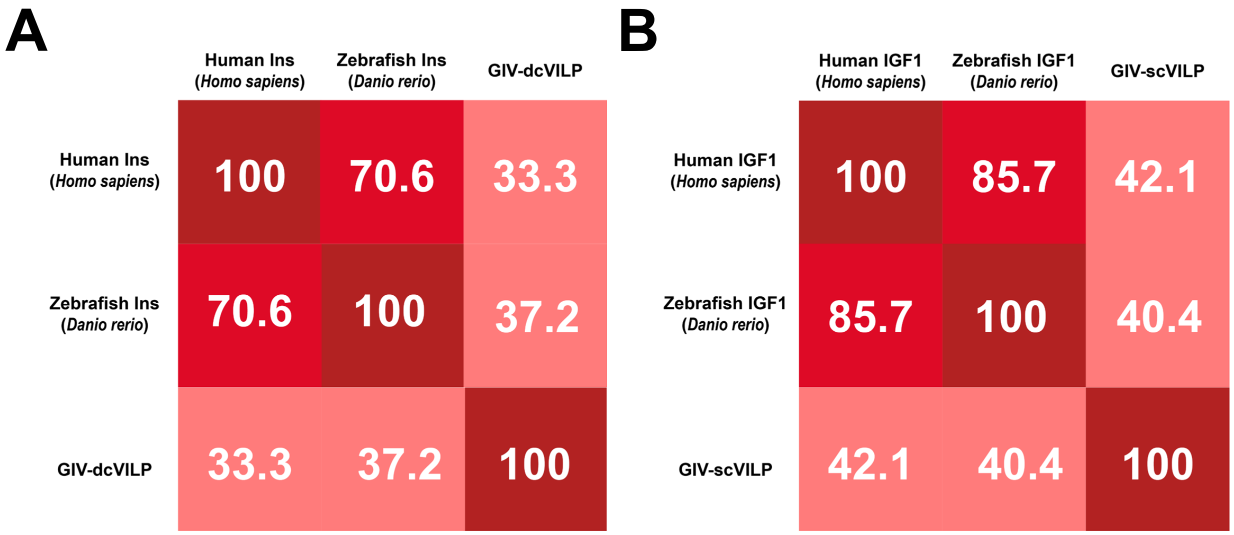


**Figure S1.** Sequence similarity among (A) double-chain peptides and (B) single-chain peptides.

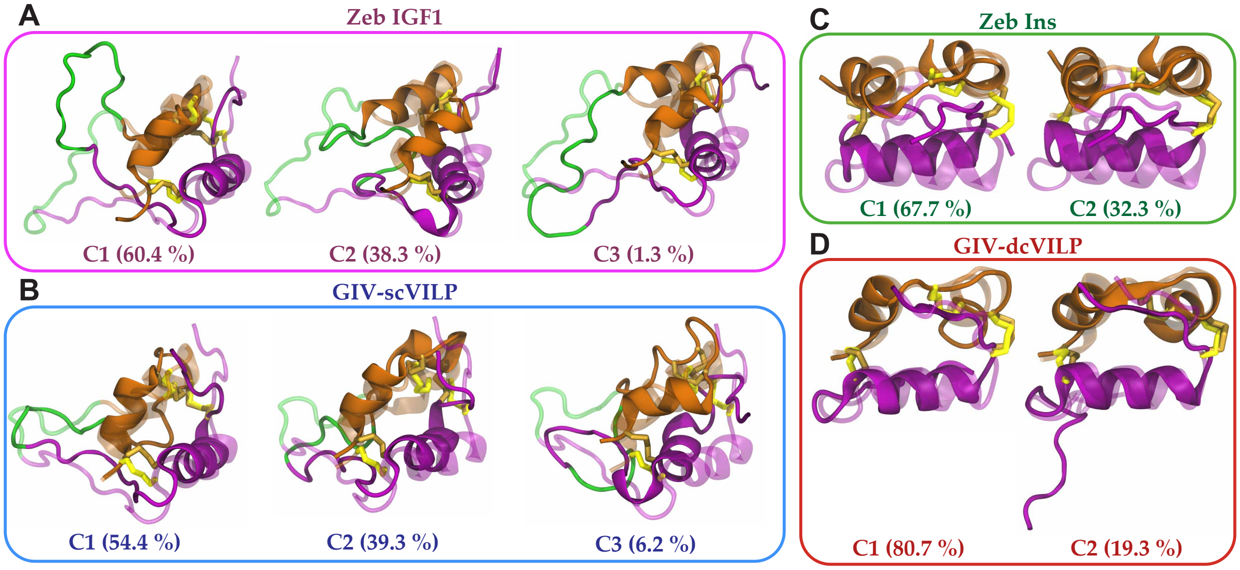


**Figure S2.** Side-view snapshots of the averaged structures (cartoon; darker colors) for each cluster of (A) Zeb IGF1, (B) GIV-scVILP, (C) Zeb Ins, and (D) GIV-dcVILP superimposed on their respective initial structures (lighter colors). cf. Figure 1 for the color scheme of peptides.

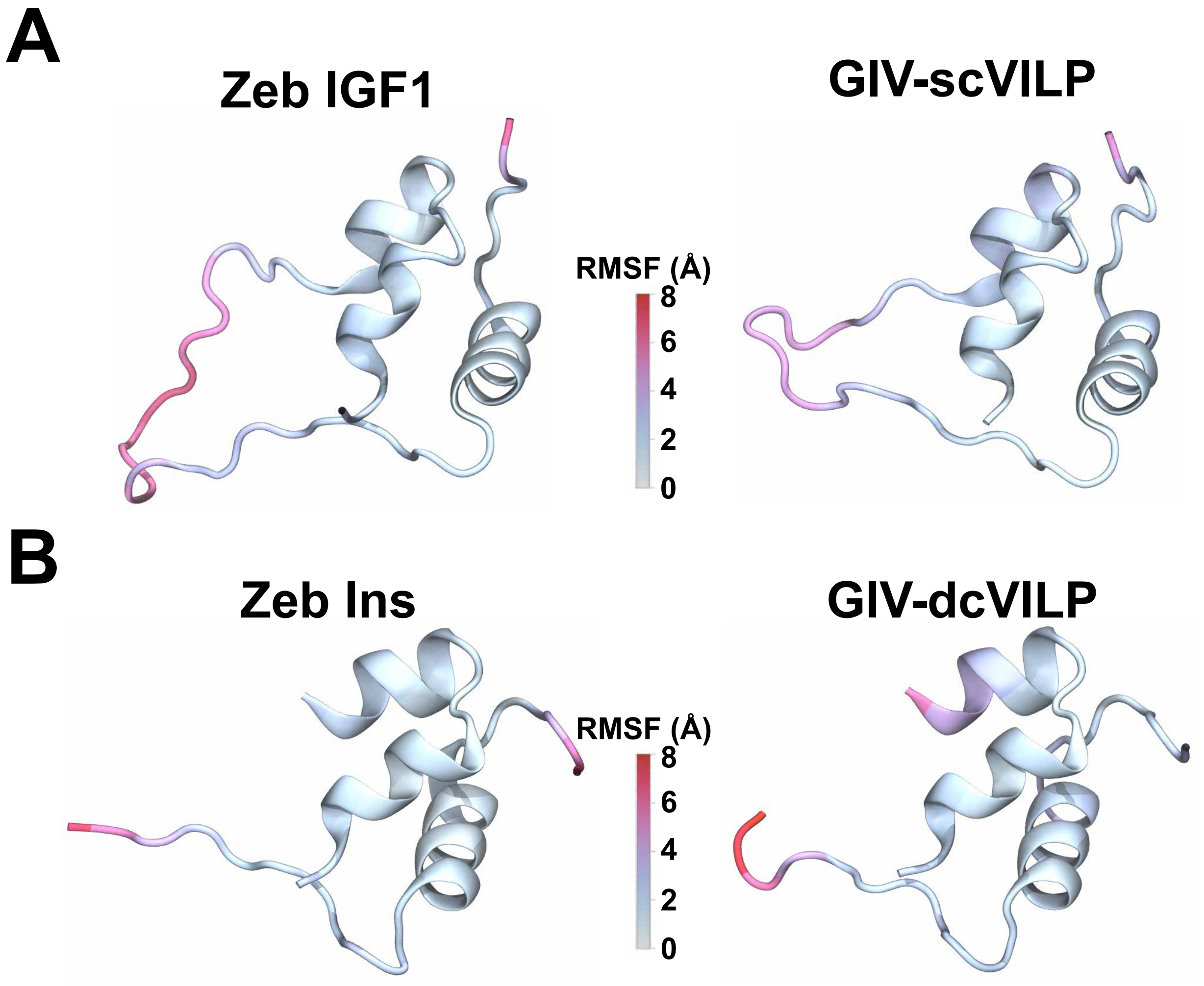


**Figure S3.** Backbone RMSF values are mapped on the structures of (A) single-chain peptides and (B) double-chain peptides. Data are based on simulations of peptides in their unbound forms.

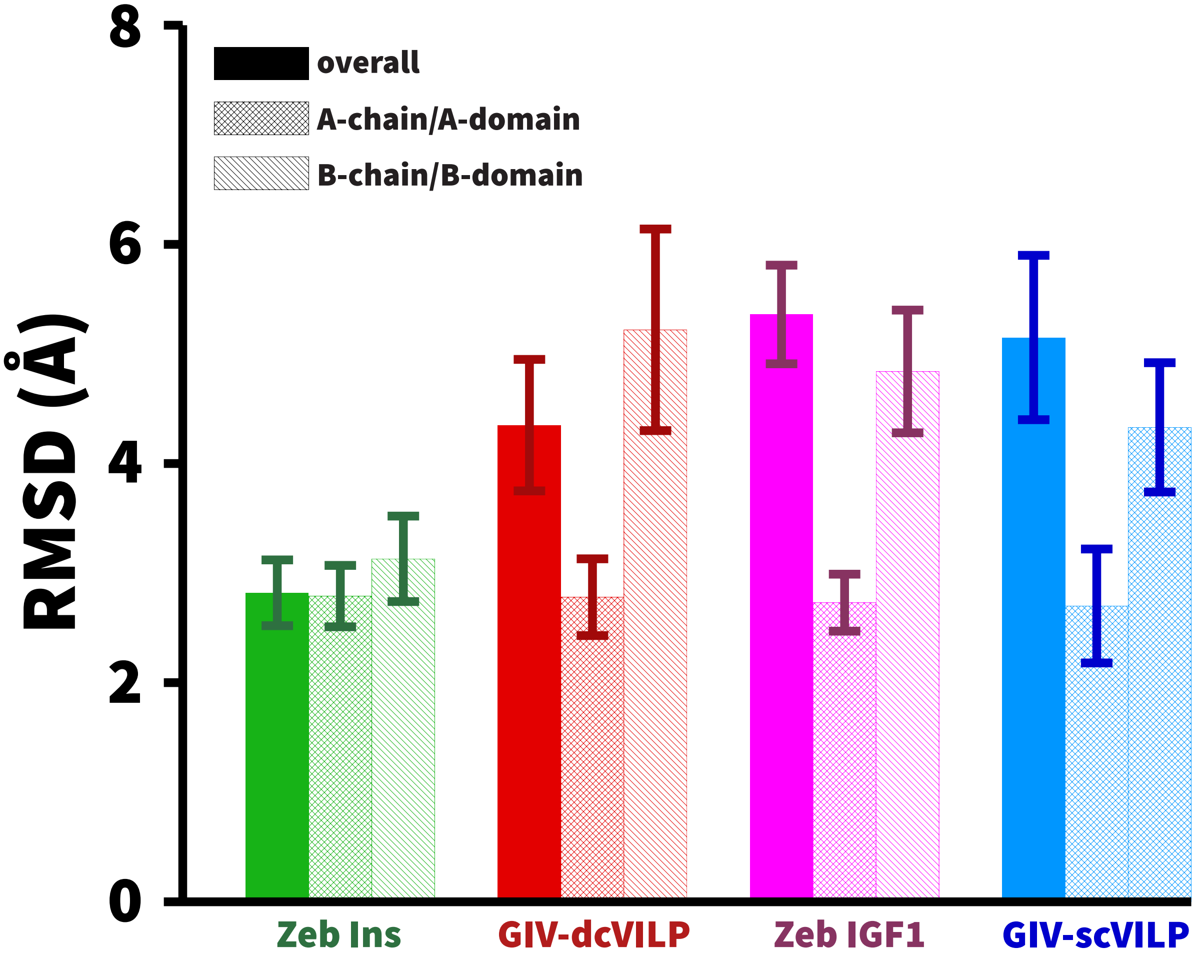


**Figure S4.** The mean RMSD values of heavy-atoms computed relative to their initial structures are shown along with corresponding standard deviations. The RMSD values were computed for the entire peptides and further decomposed into contributions from individual structural segments: A- and B-chain for Zeb Ins and GIV-dcVILP; A- and B-domain for Zeb IGF1 and GIV-scVILP.

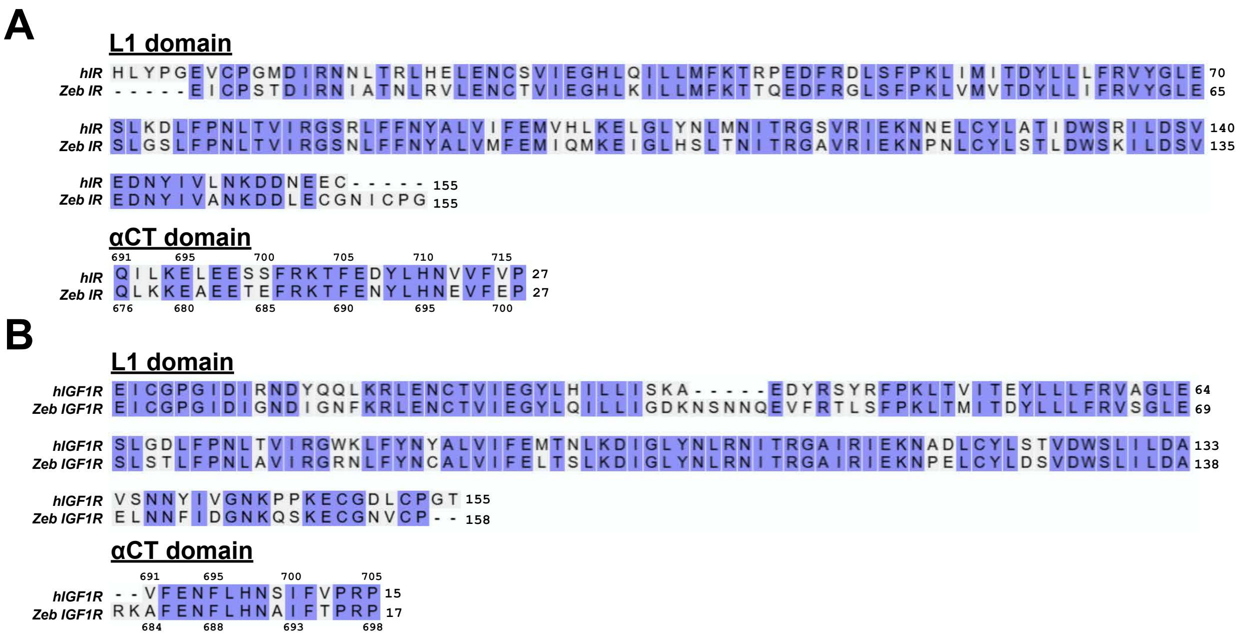


**Figure S5.** Sequence alignment of the L1 domains and the $\alpha$CT peptides between (A) human $\mu$IR (UniProtKB: P06213) and Zeb $\mu$IR (UniProtKB: Q1LVG4) and (B) human $\mu$IGF1R (UniProtKB: P08069) and Zeb $\mu$IGF1R (UniProtKB: A0A8M1PBN6) with conserved residues enclosed in blue boxes.

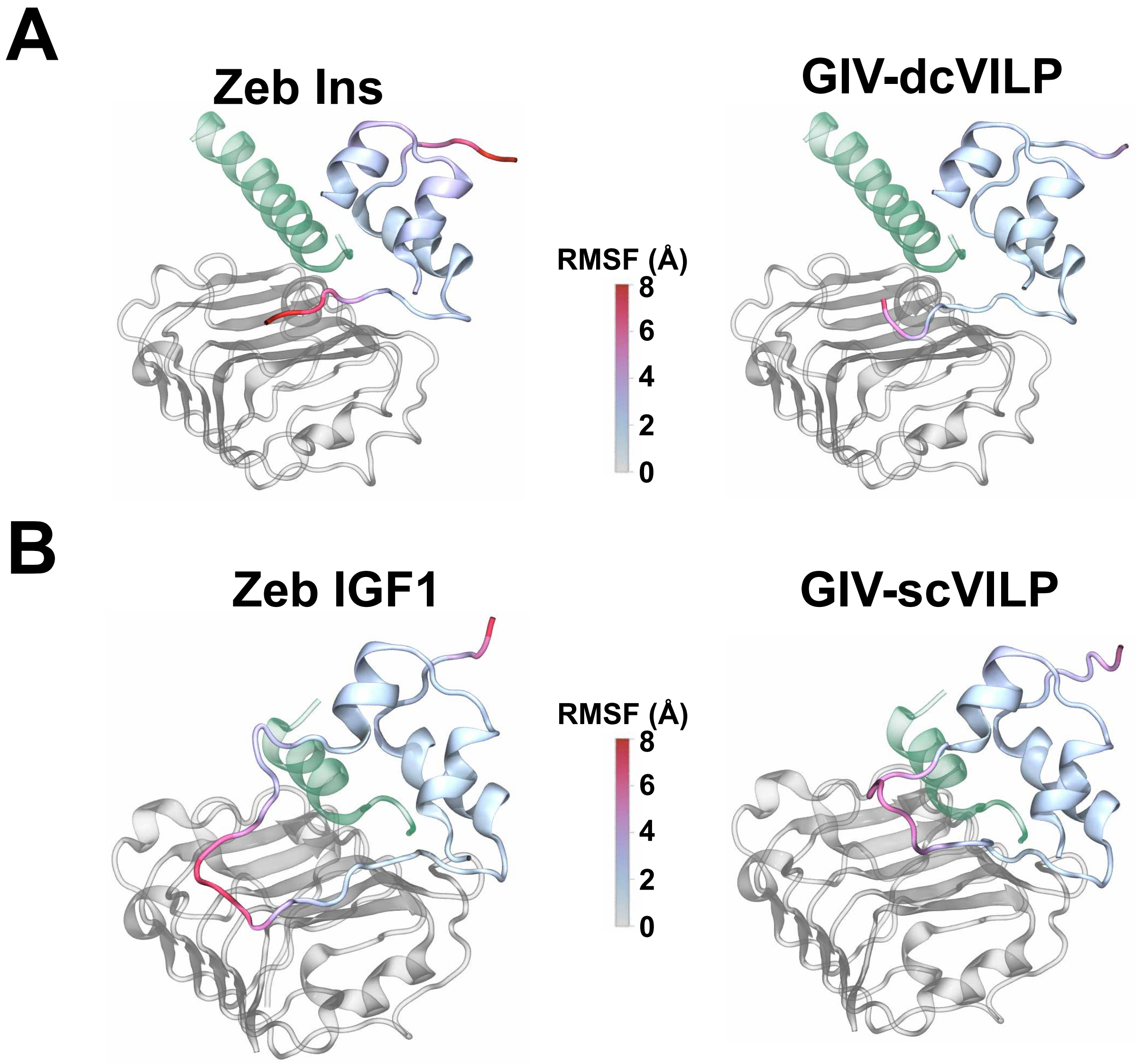


**Figure S6.** The backbone RMSF values are mapped on the structures (cartoon) of (A) double-chain peptides and (B) single-chain peptides. In each panel, the L1 domain (gray) and the $\alpha$CT peptide (dark green) are shown in a cartoon representation. Data are based on simulations of peptides in their bound forms.

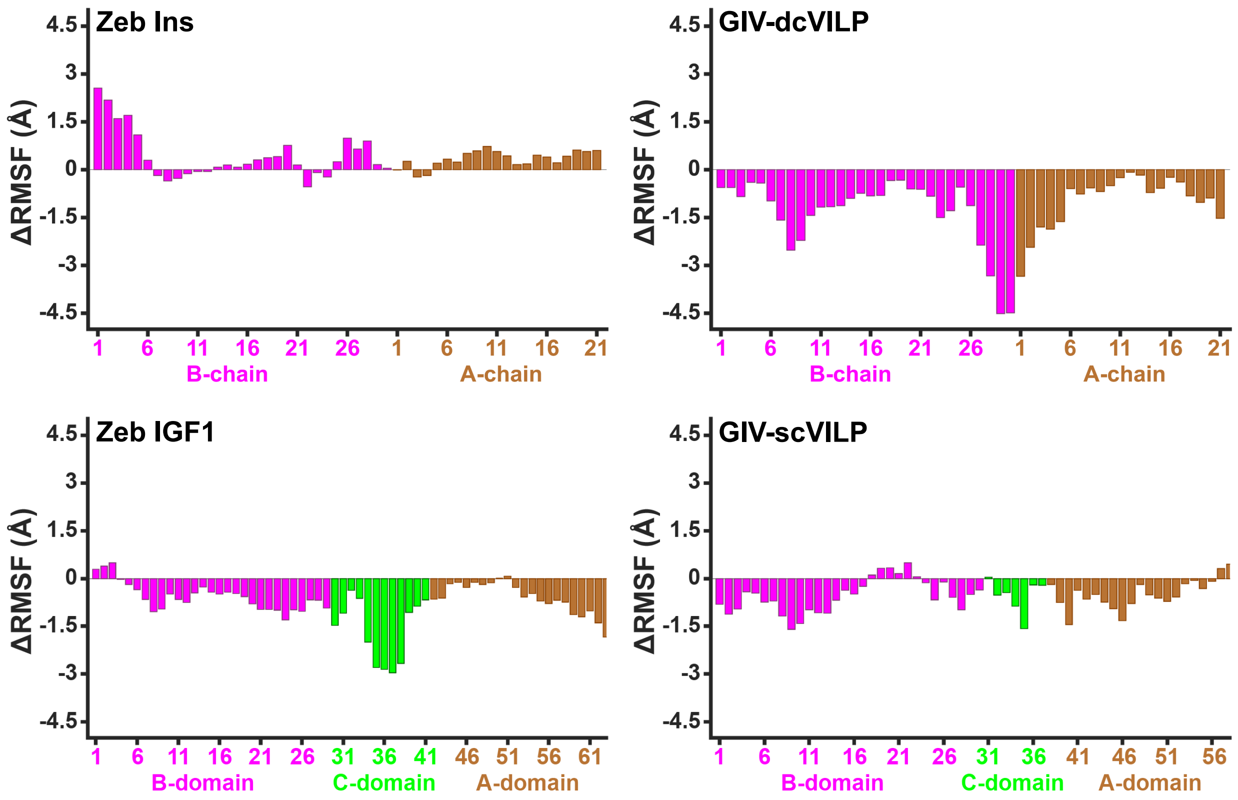


**Figure S7.** The per-residue change in root mean squared fluctuations (ΔRMSF) data highlighting the RMSF differences between the bound and unbound peptide simulations. A negative ΔRMSF value indicates that a given residue in the peptide is more stable in the bound state relative to the unbound state. The ΔRMSF values of the residues from the A-chain and the B-chain of the double-chain peptides are represented in orange and magenta bars, respectively. For the single-chain peptides, the residues from the A-domain, the B-domain, and the C-domain of each peptide are represented in orange, magenta, and green bars, respectively.

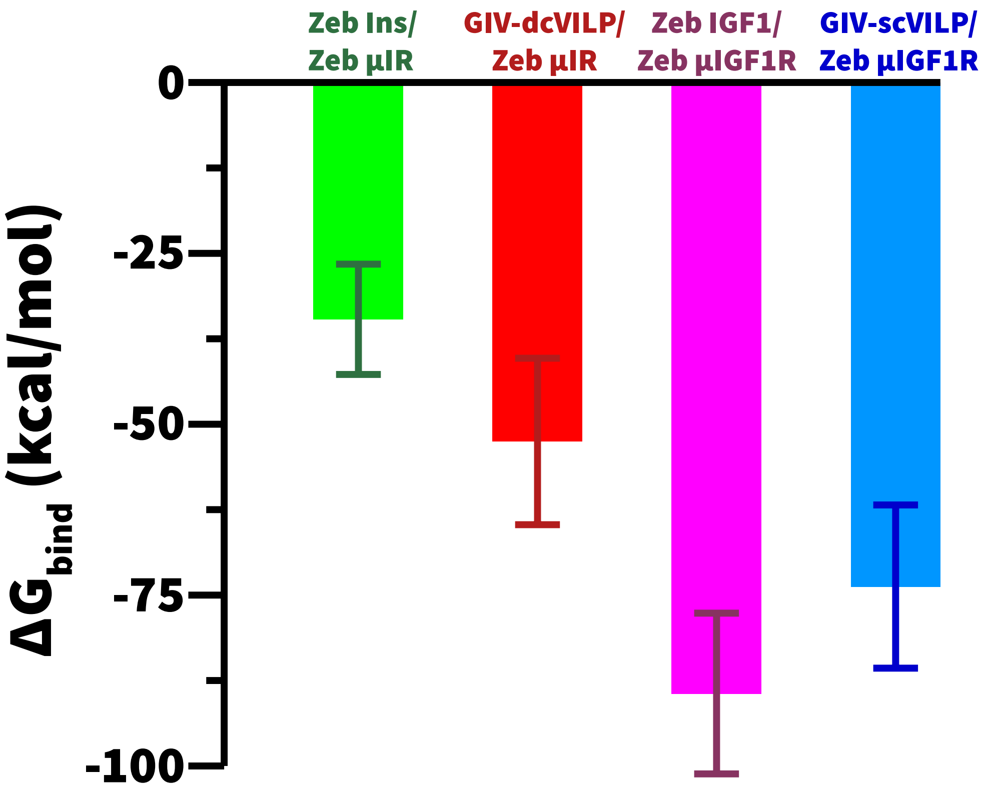


**Figure S8.** The averaged total $\Delta G_{\text{bind}}$ computed between all-atoms of each peptide ligand and of the corresponding receptor. cf. Table S4 for data.
